# Bulk delivery of a preassembled apical surface initiates epithelial lumen formation

**DOI:** 10.1101/2025.03.06.641813

**Authors:** Sujasha Ghosh, Eleanor Martin, Yuhong Chen, Natasha Amiera, Samaksh Singh, Kay En Low, Rossana Girardello, Barbara Hübner, Gunnar Dittmar, Alexander Ludwig

## Abstract

The formation of a microvilli-rich lumen is a key event in epithelial polarity development and tissue morphogenesis. During *de novo* lumenogenesis, epithelial cells establish luminal identity by directing apical cargo to an apical membrane initiation site (AMIS). Although this process has been widely studied, the mechanisms governing AMIS formation and its progression into a luminal precursor remain poorly understood. Here we combined quantitative light and electron microscopy with proximity proteomics to investigate the mechanistic basis of lumen initiation in MDCK-II cells. Contrary to prevailing models, we find that apical cargo is delivered to the AMIS in large intracellular apical precursor organelles, termed vacuolar apical compartments (VACs). VACs possess a preassembled microvilli-rich cortex and undergo exocytic fusion at the AMIS to generate a nascent lumen. Moreover, lumen initiation is tightly coordinated with the assembly and rearrangement of apical cell-cell junctions and requires the Crumbs complex protein PatJ, which controls the architecture of the apical-lateral border and connects the tight junction to the apical cortex. Together, our results identify PatJ as a critical organizer of the apical-lateral interface and indicate that VACs act as specialized transport organelles that deliver a preassembled apical cortex to the AMIS, enabling rapid and efficient lumen initiation.

## Introduction

The formation of luminal epithelial structures is critical during development and a central feature of many organs^1–6^. *De novo* lumen formation refers to the development of an apical cavity in the centre of a group of polarizing cells. Lumen initiation can be achieved through cavitation, which is mediated by apoptotic cell death, or through cord hollowing, which relies on the transport of apical determinants to an apical membrane initiation site (AMIS). Cord hollowing is utilized in the vertebrate kidney ^7–9^, the zebrafish gut ^10,11^, and the mammalian blastocyst ^12–14^. Madin-Darby Canine Kidney type II cells (MDCK-II, henceforth referred to as MDCK) also form luminal cysts via cord hollowing when cultured in Matrigel ^15^. Lumenogenesis in MDCK cells is highly stereotypic and temporally well-defined, and many mechanisms first demonstrated in this system have subsequently been shown to operate *in vivo* ^1–6,15^. MDCK cells therefore offer a powerful 3D cell culture model to study the mechanistic basis of *de novo* lumen formation.

Central to *de novo* lumenogenesis is the assembly of the AMIS and its subsequent maturation into a luminal precursor, the pre-apical patch (PAP). The AMIS is a transient and universal lumen-initiating structure ^8,9,16–18^, yet how the AMIS matures into an apical luminal surface is not clear. In MDCK cells the AMIS is formed at the cytokinetic furrow (or midbody) after the first cell division. The site of AMIS assembly is specified by specific phospholipids and polarity factors such as Par3, leading to local Rho GTPase signaling and rearrangements in the actin cytoskeleton ^16,19–21^. Apical membrane proteins (such as Crumbs-3 (Crb3) and podocalyxin (PODXL)) are then internalized from the ECM-facing cortex and delivered to the AMIS via vesicular transport. Current models imply that transcytosis of apical cargo is mediated by apical Rab GTPases including Rab11, Rab8, Rab3, and Rab35 ^16,22–25^. Once apical cargo has been delivered, the AMIS transforms into the PAP. The formation of the PAP is characterized by the assembly of tight junctions (TJs) and adherens junctions (AJs) around the nascent apical surface. In a final step, opposing apical membranes create a nascent lumen, which expands and grows in volume as cells in the cyst divide. Opening and growth of the lumen has been suggested to be mediated by electrostatic repulsion of charged apical membrane proteins (such as PODXL) and by apical proton pumps that establish electrochemical gradients leading to fluid influx into the lumen ^10,26,27^. This illustrates that *de novo* lumen formation is a hierarchical process that relies on the precise temporal coordination of membrane trafficking, cell-cell junction formation and apical domain specification and growth. However, the underlying molecular mechanisms that regulate the formation of the AMIS and its transition into the PAP are not well understood, in part because spatio-temporal and ultrastructural information of the lumen initiation process is limited.

In this study, we investigated the dynamics and ultrastructural basis of lumen initiation in MDCK cells, focusing on the epithelial Crumbs complex, a key and evolutionarily conserved regulator of epithelial tissue morphogenesis ^28,29^. The Crumbs complex comprises the transmembrane protein Crb3, the adaptor protein Pals1, and the multi-PDZ domain scaffolding protein PatJ. Crb3 interacts with Pals1 through a C-terminal PDZ-binding motif, while PatJ associates with Pals1 via its N-terminal L27 domain ^30–34^. In mice, loss of *Crb3* or *Pals1* disrupts tissue architecture in multiple organs, including the intestine, lung, and kidney ^35–37^. In MDCK cells, silencing or inactivation of Crb3, Pals1, or PatJ impairs TJ assembly and results in a multi-lumen phenotype, characterized by the formation of multiple ectopic lumens instead of a single apical lumen ^33,38–44^. However, the mechanisms by which the Crumbs complex regulates lumen formation remain unclear.

By combining quantitative light and electron microscopy with time-resolved proximity proteomics, we demonstrate that lumen initiation is temporally and spatially coordinated with the assembly of the apical junctional complex (AJC). Unexpectedly, we also discovered that apical cargo is delivered to the AMIS via large intracellular organelles known as vacuolar apical compartments (VACs), which fuse with the AMIS to initiate apical domain identity. Furthermore, we show that PatJ is essential for VAC exocytosis, and consequently, for lumen initiation.

## Results

### Pals1 and apical cargo are delivered to nascent cell-cell contacts via VACs

Using proximity labeling proteomics and transmission electron microscopy (TEM) with the peroxidase APEX2 ^45^ we previously demonstrated that the Crumbs complex forms a distinct domain apical of TJs, the so-called vertebrate marginal zone (VMZ) ^43^. The spatial separation of the Crumbs complex from apical cell junctions is evolutionarily conserved ^46^, suggesting a specific role at the apical-lateral interface. We set out to study the spatio-temporal assembly of the VMZ using a stable MDCK cell line expressing a Pals1-APEX2-EGFP (Pals1-A2E) fusion protein. The calcium switch model (Fig. 1A) was used to determine the localization of Pals1 at distinct stages of cell-cell junction formation by light microscopy and TEM. In addition, we used the Pals1-A2E protein as a proxy to study the assembly and maturation of the apical-lateral border using time-resolved proximity proteomics.

**Figure 1:**
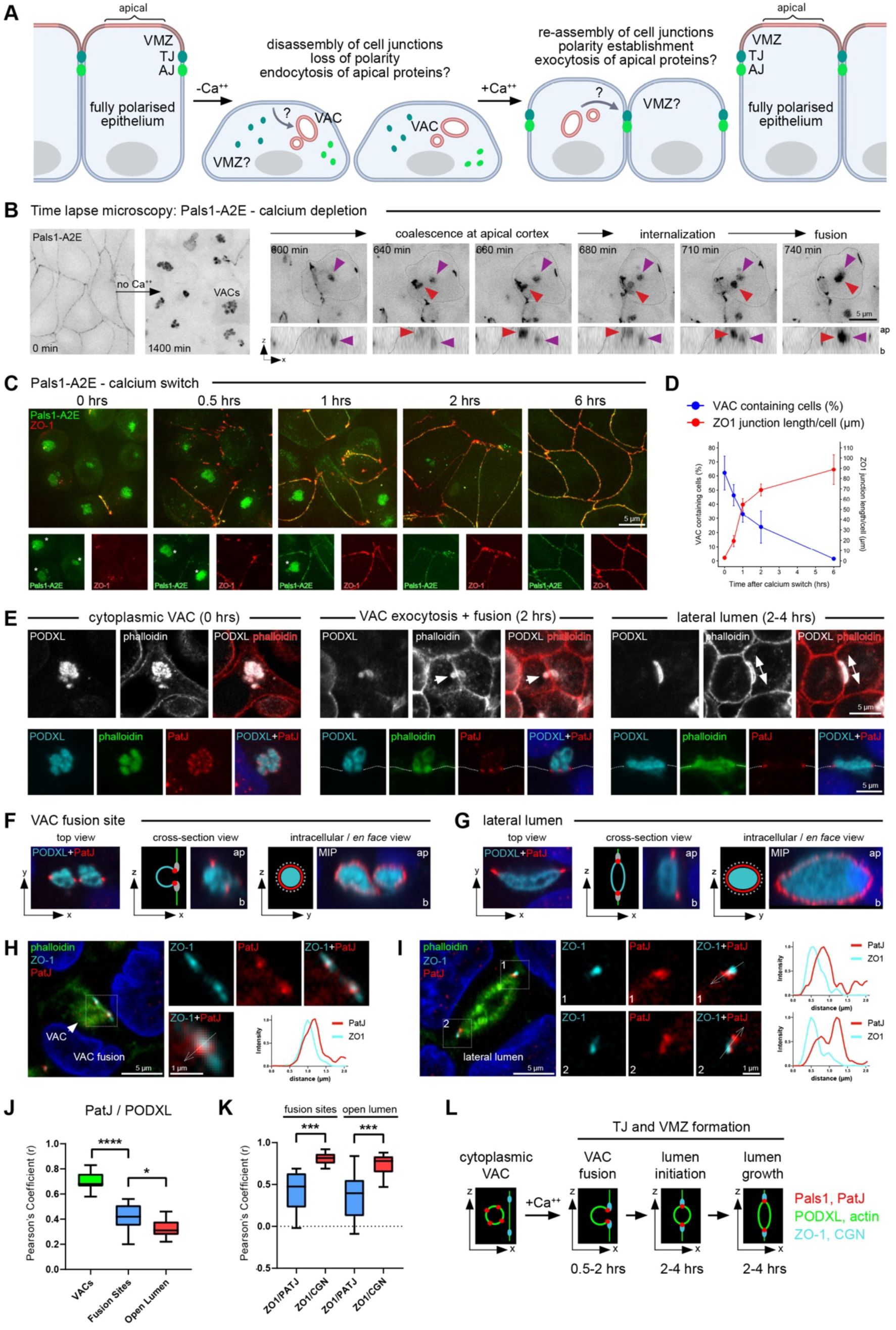
**VACs deliver apical cargo to nascent cell junctions to initiate lateral lumens** (A) Schematic illustrating the calcium switch assay. How VACs are internalized and subsequently trafficked back to the cortex is unknown. The trafficking and functions of VMZ proteins are also unclear. (B) Time-lapse imaging of MDCK cells stably expressing Pals1-A2E upon calcium withdrawal. Pals1 is internalized into large organelles (VACs) in response to calcium depletion. Red and purple arrowheads demarcate a VAC internalization event and an existing intracellular VAC, respectively. VAC fusion events are observed at 740 min. (C) Calcium switch assay in Pals1-A2E cells. Cells were fixed at the indicated time points post-calcium addition and stained with antibodies against ZO-1. See also Figure S1. (D) Quantification of tight junction length and VAC frequency based on the data shown in (C). Data is expressed as mean +/- SD (n=115-167 for each time point). (E) MDCK cells were fixed 0-4h post calcium addition and stained with antibodies against PODXL and PatJ, and phalloidin to stain F-actin. Intracellular VACs (left), VAC fusion sites (middle), and lateral lumens (right) are shown. (F and G) Cross-sections and projection images of VAC fusion sites (F) and lateral lumens (G). MDCK cells were fixed at 2h post calcium addition and stained with anti-PatJ and anti-PODXL antibodies. (H and I) VAC fusion sites (H) and lateral lumens (I) stained with antibodies against ZO-1 and PatJ, and phalloidin. Single confocal sections and line scans along the junctional-luminal interface are shown. (J and K) Pearson’s correlation coefficients of PODXL, TJ and VMZ markers in VACs, at fusion sites, and at lateral lumens. * p<0.05, *** p<0.001; **** p<0.0001, Student’s t-test. (n=10-15). See also Figure S2. (L) Model of VAC fusion and lateral lumen formation.

In confluent MDCK monolayers, Pals1-A2E localized to apical cell junctions, as expected (Fig. 1B). Following overnight calcium depletion Pals1 had redistributed from the cortex into large intracellular structures reminiscent of previously described vacuolar apical compartments (VACs) ^47,48^ (Fig. 1B). Time-lapse microscopy revealed that loss of extracellular calcium led to the progressive dissolution of cell-cell junctions and loss of polarity within 6h. This was followed by the coalescence of Pals1 at the apical (dorsal) pole and its internalization into VAC-like structures, which appeared to fuse or coalesce within the cytoplasm (Fig. 1B). To determine whether VACs contained endocytosed apical membrane components, we applied a fluorescently labeled antibody against the apical glycoprotein PODXL to live monolayers prior to calcium removal. PODXL and Pals1 clustered at the cell cortex and were subsequently co-internalized into VACs (Fig. S1A). Consistent with their apical origin, VACs also contained the apical protein ezrin and additional VMZ proteins (PatJ and Homer proteins), whereas TJ proteins (ZO-1 and Par3) were not present in VACs and instead appeared as small cytoplasmic puncta (Fig. S1B). Moreover, VACs were enriched in F-actin, contained the apical calcium pump PMCA2, and were stained by the calcium indicator Fluo-4 (Fig. S1B-S1D). Calcium depletion in Caco-2 monolayers also resulted in the appearance of VAC-like structures, indicating that their formation is not restricted to MDCK cells (Fig. S1E, S1F). We concluded that loss of polarity causes VMZ and apical proteins to be internalized into an intracellular organelle characteristic of VACs ^47,48^, or apicosomes ^49^.

When calcium-depleted cells were shifted back to normal calcium media, VACs progressively disappeared and were no longer detectable 6h post calcium replenishment (Fig. 1C). At the same time, TJs reformed, resulting in a continuous junctional belt containing both ZO-1 and Pals1-A2E. Quantification revealed an inverse relationship between VAC frequency and TJ length during junction re-establishment, indicating that VAC exocytosis and TJ formation are temporally coordinated (Fig. 1D). To understand when and where VACs are exocytosed, we initially analysed the localization of PODXL, F-actin (stained with phalloidin) and PatJ (as a VMZ marker) in wild-type MDCK cells at different stages of the calcium switch assay (Fig. 1E, Fig. S2A-S2D). In calcium-depleted cells (0h), PatJ, PODXL and F-actin colocalized in VACs, as expected (Fig. 1E, left panel). Interestingly, at 2h post calcium addition, apparent VAC exocytosis and nascent VAC fusion events with the lateral membrane were evident (Fig. 1E, middle panel). At this stage, PatJ formed a distinct belt around VAC fusion sites (Fig. 1F), indicating that it had translocated into the lateral membrane. At slightly later time points (2-4h), elongated, F-actin rich and PODXL-containing pockets were observed along the lateral membrane (Fig. 1E, right panel). These pockets displayed an apparent paracellular space and were again surrounded by a belt containing PatJ (Fig. 1G).

The observations above prompted us to quantify the relative co-localization of VMZ proteins (Pals1 and PatJ), TJ proteins (ZO-1 and cingulin (CGN)), and apical proteins (PODXL, F-actin) during junction re-establishment using Airyscan microscopy (which offers a lateral resolution of ∼120 nm) (Fig. 1H-1K, Fig. S2E-S2G). We found that both Pals1 and PatJ colocalized efficiently with PODXL or F-actin in intracellular VACs and that colocalization decreased significantly at the “VAC fusion” and “open lumen” stages (Fig. 1J, Fig. S2F, S2G). Interestingly, although both VMZ and TJ proteins formed a junctional ring around VAC fusion sites and around lateral lumens (Fig. 1H, 1I, Fig. S2E), TJ proteins (ZO-1/CGN) colocalized significantly more strongly with each other than with PatJ or Pals1 (Fig. 1K, Fig. S2E-S2G). Consistent with this, PatJ and Pals1 were invariably located closer to the apical cortex than ZO-1 or CGN (Fig. 1H, 1I). This suggests that even at these early stages of junction formation, VMZ and TJ proteins are spatially separated. We concluded that VACs deliver apical cargo to nascent cell-cell contacts to initiate the assembly of a lateral luminal space. Such lateral lumina are surrounded by a junctional belt composed of distinct VMZ and TJ domains (Fig. 1L).

### VACs are exocytosed at nascent cell-cell contacts

Next, we analysed the ultrastructure of VACs and their sub-cellular redistribution during cell-cell junction assembly using APEX2-TEM. Pals1-A2E MDCK cells were fixed at 0h, 2h, and 6h of the calcium switch assay and subjected to APEX2 labeling (Fig. 2A). The APEX2 tag serves as an electron-dense label, which aided in the identification of VACs in TEM and allowed us to localise Pals1 within VACs with 5-10 nm precision ^50^. In calcium-depleted cells, VACs appeared as large, micrometer-size intracellular vacuoles rich in apical microvilli (Fig. 2B (0h), Fig. S3). Microvilli often appeared more electron dense than the surrounding membranes, indicating the presence of Pals1 in these structures (Fig. 2B (0h), black arrowheads, Fig. S3A). VACs were frequently surrounded by small vesicles (Fig. 2B (0h), white arrowheads, Fig. S3A and S3B). At 2h post calcium addition, fusion events of VACs with the lateral membrane were evident (Fig. 2B (2h)), with the fusion pore being clearly visible. In addition, elongated microvilli-rich lumens were observed at the lateral domain (Fig. 2B (2h)). Interestingly, Pals1 APEX2 label was observed at cell-cell contacts immediately adjacent to the VAC fusion pores and lateral lumens (Fig. 2B (2h), Fig. S4A). These TEM micrographs correlate with the data shown in Figure 1E-1G and show that VACs indeed fuse at nascent cell-cell junctions.

**Figure 2:**
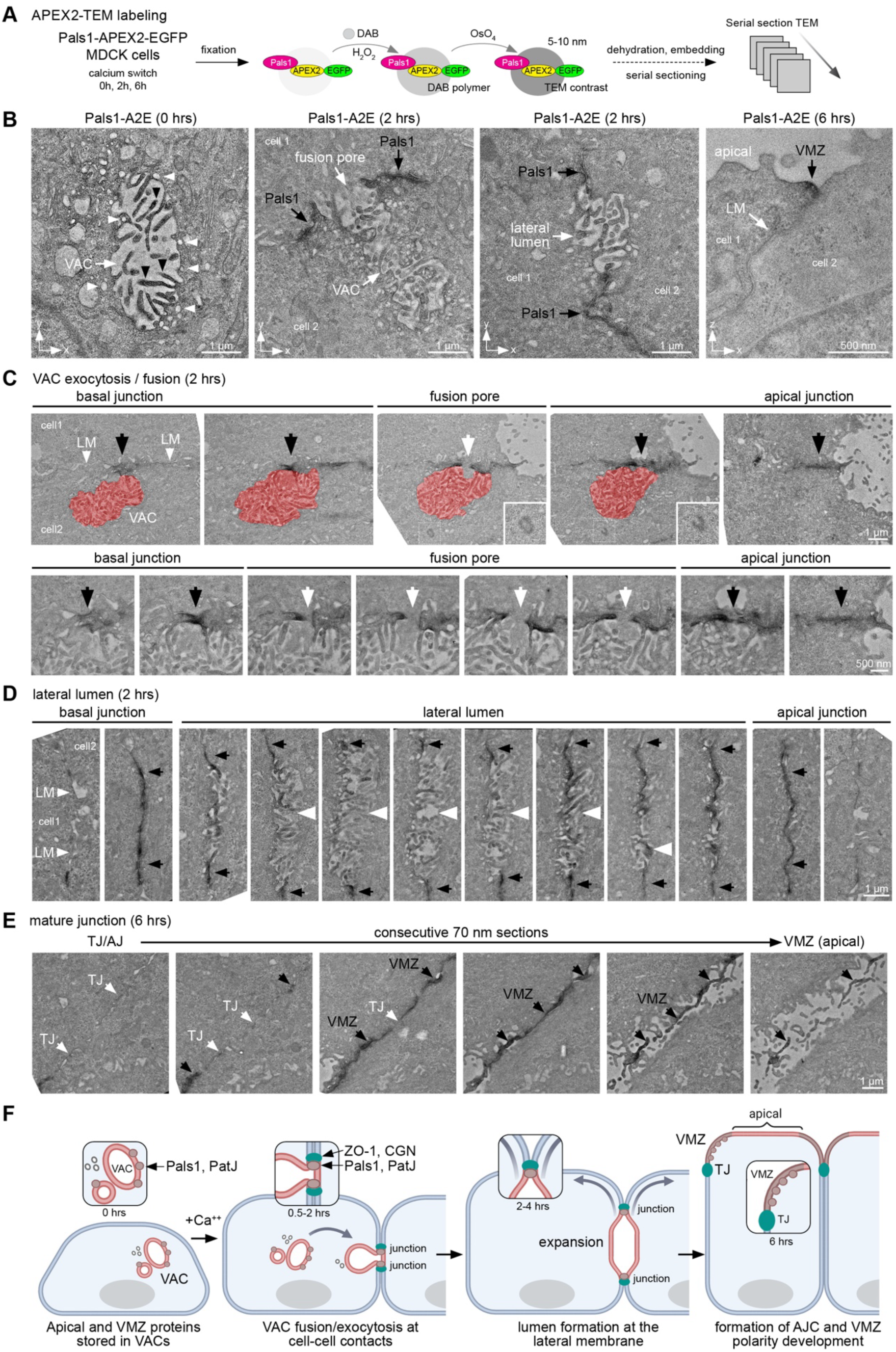
**Ultrastructural analysis of VAC exocytosis and lateral lumen formation** (A) Principle of APEX2-TEM labeling. A brief (2-5 min) incubation of fixed cells with diaminobenzidine (DAB) and hydrogen peroxide (H_2_O_2_) results in the deposition of a DAB polymer that upon binding to OsO_4_ produces an electron-dense stain. (B) APEX2 TEM labeling of Pals1-A2E at different time points of the calcium switch assay. White arrowheads indicate small vesicles close to VACs, black arrowheads indicate enhanced contrast of microvilli inside the VACs produced by APEX2 labeling. LM = lateral membrane. See also Figure S3. (C) Serial section TEM of a VAC fusion event with the lateral membrane. Select serial sections of the volume are shown. Note the increase in junctional EM contrast produced by Pals1-A2E (black arrowheads) around the VAC fusion site (white arrowhead). Centrioles are localised close to the VAC fusion site (insets). See also Figure S4. (D) Representative serial section TEM of a lateral lumen. Selected serial sections of the volume are shown. Note the increase in junctional EM contrast produced by Pals1-A2E (black arrowheads) around the lumen (white arrowhead). See also Figure S4. (E) Representative serial section TEM of Pals1-A2E at 6h post-calcium addition. Samples were sectioned *en face* (parallel to the substratum). Consecutive serial sections of the apical/lateral interface are shown. Note the increase in EM contrast produced by Pals1-A2E (black arrowheads) apical of TJs (white arrowheads). (F) Model of lateral lumen formation and polarity development in the calcium switch assay.

To analyse the ultrastructure of VAC fusion sites and lateral lumens in more detail, we performed a comprehensive serial section TEM analysis of Pals1-A2E MDCK cells 2h post calcium addition. Embedded cell monolayers were sectioned parallel to the growth support, producing consecutive *en face* TEM sections spanning ∼5 µm of cell volume (Fig. 2A). Serial section TEM of VAC fusion sites revealed a strong electron-dense label above (apical), below (basal), and adjacent to the fusion pore (Fig. 2C, Fig. S4B-D). This confirms that upon fusion of VACs with the lateral membrane, Pals1 rapidly accumulates at nascent cell-cell contacts. Lateral lumens rich in microvilli were also surrounded by cell-cell junctions labeled by Pals1. Labeling was again evident both apical and basal of the lumen opening (Fig. 2D, Fig. S4E). Interestingly, centrioles were frequently observed close to VAC fusion sites (Fig. 2C, Fig. S4C, insets) and were also often associated with intracellular VACs (Fig. S3C), suggesting a role for the centrosome in VAC trafficking or membrane targeting. Lastly, serial section TEM of *en face* sections 6h post calcium addition (a time point at which cell junctions are fully assembled) revealed Pals1 APEX2 labeling associated with microvilli-like protrusions just apical of cell-cell junctions (Fig. 2E). TEM of monolayers imaged in cross-sections confirmed this apical localization of Pals1 in polarized cells (Fig. 2B (6h)). We concluded that VAC exocytosis is coordinated with the formation of cell-cell junctions and results in the emergence of an apical luminal surface at the lateral membrane (Fig. 2F).

### A dynamic protein network governs VAC exocytosis and cell-cell junction assembly

We wanted to understand the molecular basis of the complex ultrastructural rearrangements that occur during the assembly and maturation of the AJC. To this end, we generated proximity proteomes of Pals1-A2E at distinct stages of the calcium switch assay using previously established protocols ^43,51^ (Fig. 3A). Principal Component Analysis showed a strong correlation between replicates of the same time point, and early and late time points were clearly separated from each other (Fig. S5A). After removal of common contaminants and proteins that were not significantly enriched with Pals1 compared to the controls, we arrived at a proteome of 278 proteins (Fig. S5B). The 278 proteins dataset was subjected to unsupervised hierarchical clustering, resulting in four clusters with distinct kinetics (Fig. 3B, 3C, Fig. S5C and S5D). Cluster 4 contained proteins that were rapidly depleted from the Pals1 proteome upon calcium addition, cluster 1 and cluster 2 contained proteins that became transiently enriched at 0.5h and 2h, respectively, and cluster 3 contained proteins whose log2 fold changes progressively increased, reaching peak enrichment at 6h (Fig. 3B, 3C). Gene Ontology analysis showed that clusters 1 and 2 were enriched in proteins associated with protein transport and GTPase signaling, whilst cluster 3 was enriched in proteins involved in cell-cell adhesion and the regulation of the actin cytoskeleton (Fig. S5E). Moreover, cluster 3 was enriched in proteins previously identified in a static Pals1 proximity proteome determined from fully polarized MDCK cell monolayers (Fig. S5F) ^43^.

**Figure 3:**
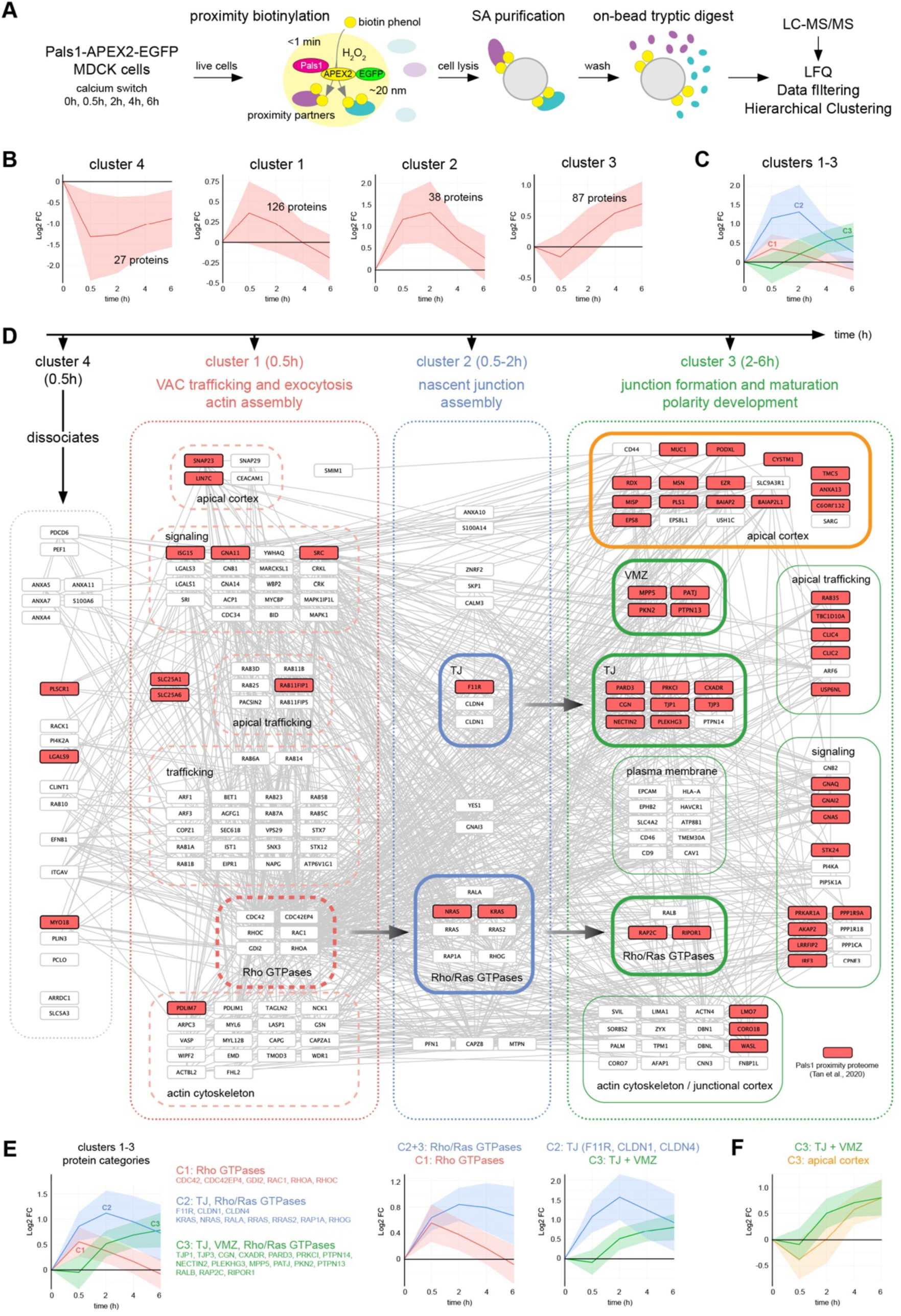
**Time-resolved proximity proteomics resolves distinct stages of VAC trafficking and cell-cell junction assembly** (A) Principle of proximity biotinylation with APEX2 and workflow used to generate a time-resolved proteome of Pals1 during cell-cell junction assembly in the calcium switch assay. Live cells expressing Pals1-A2E are incubated with biotin phenol for 30 min followed by the addition of H_2_O_2_ for 1 min. This results in the biotinylation of proteins in a radius of ∼20 nm. Upon cell lysis, biotinylated proteins are isolated using streptavidin (SA) pulldowns and analysed by liquid chromatography tandem mass spectrometry (LC-MS/MS). (B) Hierarchical clustering of the 278 identified proteins reveals four clusters with distinct kinetics. Data is expressed as mean +/- SD. (C) Direct comparison of the kinetics of clusters 1-3. (D) Temporally resolved protein interaction network of Pals1 during cell-cell junction assembly based on STRING and BioGRID entries. A total of 200 proteins (out of 278 identified) are shown. Proteins were categorized based on known functions and our previously defined proteome of the apical-lateral border ^43^. Proteins highlighted in red are core components of the static Pals1 proteome identified in Tan et al., ^43^. See also Figure S5. (E) Kinetics of Rho and Ras GTPases, TJ proteins and VMZ proteins. (F) Kinetics of TJ and VMZ proteins and proteins residing at the apical cortex.

To visualise the molecular composition, connectivity and temporal evolution of the Pals1-associated protein network, we generated a temporally resolved protein-protein interaction network based on STRING and BioGRID database entries (Fig. 3D). For simplicity, only 200 out of the 278 proteins identified were used for this analysis (cytosolic, ribosomal, and miscellaneous proteins were not considered). Proteins were grouped according to established functions and based on our previous proteomics analysis of the apical-lateral border ^43^. Cluster 4 contained proteins involved in calcium binding and signaling, suggesting that such proteins dissociate from VACs soon after extracellular calcium replenishment. Cluster 1 contained all major Rho-type GTPases (RhoA, RhoC, Rac1, and Cdc42) and regulators of the actin cytoskeleton including actin assembly factors and myosin-II motor subunits. In addition, Rab and SNARE proteins involved in apical trafficking (Rab11B, Rab25, Rab3D) and endosomal transport (Rab5B, Rab5C, SNX3, STX12, STX7) were present in cluster 1. Light microscopy and TEM of APEX2 fusion proteins showed that Rab3, Rab11, and Rab25 are associated with vesicles surrounding the VACs, whereas Rab35 specifically labeled microvilli facing the VAC lumen (Fig. S6). This validates our proteomics data and suggests a role of these Rabs in VAC assembly or trafficking. Cluster 2 encompassed proteins that became enriched with Pals1 at 0.5-2h. Unlike proteins in cluster 1 (which peaked at 0.5h and became de-enriched after 4h), those in cluster 2 remained associated with Pals1 until 6h of the calcium switch. Interestingly, this cluster contained integral membrane proteins of the TJ (F11R (JAM-A), CLDN1 (claudin-1) and CLDN4 (claudin-4)) as well as RhoG, Rap1A, and additional Ras-type GTPases. We then examined cluster 3, i.e. proteins that were highly enriched at late stages of the calcium switch (at 2-6h). This cluster contained VMZ components (MPP5/Pals1, PATJ, PKN2 and PTPN13), TJ scaffolding proteins (TJP1 (ZO-1), TJP3 (ZO-3), PARD3 (Par3), PRKCI (atypical PKC), CGN (cingulin)), apical membrane proteins (PODXL, MUC1, CD44), and the ERM proteins EZR (ezrin), RDX (radixin), and MSN (moesin). In addition, Rap2C and components of the junctional actin cortex were enriched at mature cell junctions. Figure 3E illustrates the differential kinetics of Rho and Ras GTPases and the stepwise recruitment of TJ and VMZ proteins. Interestingly, closer examination of cluster 3 revealed that apical proteins become enriched with Pals1 much later than junctional proteins (Fig. 3F), suggesting that the physical linkage between the VMZ and the apical cortex occurs after the AJC is fully assembled. We concluded that the assembly and maturation of the AJC is a hierarchical process that is driven by extensive molecular remodeling of the junctional proteome. In addition, the temporally resolved protein interaction network mirrors the subcellular dynamics of Pals1 during cell junction assembly visualized by TEM.

### Dynamics of TJ and VMZ proteins during lumen initiation in MDCK 3D cultures

Our data from the calcium switch assay indicated that VAC exocytosis at nascent cell contacts leads to the formation of a lateral lumen surrounded by cell-cell junctions. To study this process in a more physiological context, we analysed lumen initiation in the MDCK 3D cyst assay (Fig. 4A, see introduction). In mature cysts PatJ and ZO-1 localised to apical cell junctions, whilst ezrin labeled the apical cortex (Fig. 4B, 4C). Airyscan microscopy and quantitative line scan analysis revealed that PatJ and ZO-1 were offset by ∼150 nm, with PatJ localising more apical than ZO-1 (Fig. 4C, 4D). This agrees with previous work and demonstrates that the VMZ is a characteristic feature of the MDCK 3D cyst system ^42,43,52^.

**Figure 4:**
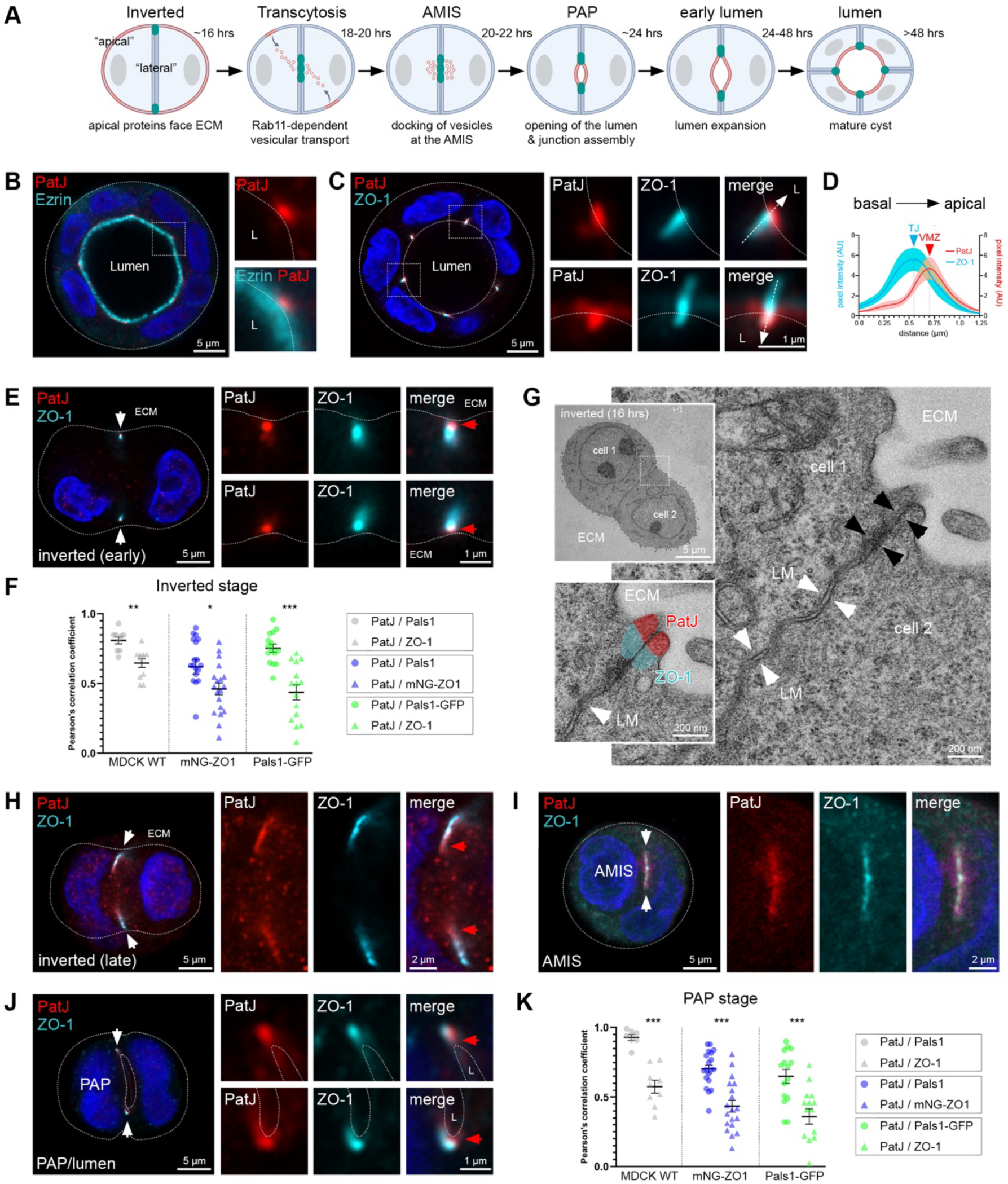
**Dynamics of TJ and VMZ proteins during lumen initiation in 3D cultures** (A) Current model of *de novo* lumen formation in MDCK 3D cultures grown in Matrigel. (B and C) Representative Airyscan images of MDCK cysts grown in Matrigel for 4 days stained with anti-PatJ and anti-ezrin antibodies (B) or with anti-PatJ and anti-ZO-1 antibodies (C). Note that PatJ localizes apical of ZO-1 (insets in C). (D) Line scan quantification of ZO-1/PatJ junctional distribution as shown in (C). Data is expressed as mean ± SD (n=10). (E) Representative (early) inverted stage. Cells were fixed at 16h in Matrigel and costained with anti-PatJ and anti-ZO-1 antibodies. (F) Pearson’s correlation coefficients of PatJ, Pals1, and ZO-1 in the inverted stage in MDCK WT cells, mNeonGreen-ZO1, and Pals1-GFP cells. * p<0.05, ** p<0.01, *** p<0.001, Student’s t-test (n=10-20). See also Figure S7. (G) Representative TEM micrographs of the inverted stage. The inset at the bottom left shows the putative localisation of PatJ and ZO-1 at the ECM-facing cell junction. (H) Representative (late) inverted stage. Cells were fixed at 16h in Matrigel and costained with anti-PatJ and anti-ZO-1 antibodies. (I) Representative AMIS stage. Cells were fixed at 20h in Matrigel and costained with anti-PatJ and anti-ZO-1 antibodies. (J) Representative PAP stage. Cells were fixed at 24h in Matrigel and costained with anti-PatJ and anti-ZO-1 antibodies. (K) Pearson’s correlation coefficients of PatJ, Pals1, and ZO-1 in the PAP stage in MDCK WT cells, mNeonGreen-ZO1, and Pals1-GFP cells. *** p<0.001, Student’s t-test (n=10-22). See also Figure S7.

To understand how and when this junctional compartmentalization is established, we analysed the localization of VMZ and TJ proteins at early and late 2-cell stages using Airyscan microscopy (Fig. 4E-4K, Fig. S7). At early time points (inverted stage; 16-18h), PatJ, Pals1, and ZO-1 formed two discrete puncta at the periphery of the cell-cell contact interface facing the ECM (Fig. 4E, Fig. S7B). Notably, although PatJ/Pals1 and ZO-1 fluorescence overlapped, their distributions were not fully congruent: Whilst PatJ and Pals1 were positioned toward the ECM-facing side, ZO-1 extended toward the center of the cell-cell interface. Consistent with this observation, Pearson’s correlation analysis showed that PatJ colocalized significantly more strongly with Pals1 than with ZO-1 (Fig. 4F). Interestingly, TEM of early-stage cell doublets revealed close membrane appositions at the periphery of the cell-cell contact interface, consistent with the presence of TJs (Fig. 4G, black arrowheads). By contrast, lateral cell-cell contacts showed a wider intermembrane space and frequently appeared highly interdigitated (Fig. 4G, white arrowheads, Fig. S8), likely reflecting the presence of E-cadherin-based adhesions ^53^. This suggests that prior to lumen initiation, TJs and the VMZ form an “inverted” junctional complex that separates the “apical” ECM-facing cortex from the lateral membrane interface (Fig. 4G).

In addition to inverted stages with a clear “double puncta” pattern (Fig. 4E), we also observed stages in which Pals1/PatJ and ZO-1 appeared to “spread along” the lateral membrane interface (Fig. 4H, Fig. S7C). Interestingly, Pals1/PatJ were now oriented toward the center of the cell doublet, whereas ZO-1 was positioned more peripherally. This indicates a spatial inversion in the organization of VMZ and TJ components relative to the inverted stage. At slightly later time points (20-22h), TJ and VMZ proteins coalesced at the center of the lateral membrane interface, demarcating the AMIS (Fig. 4I, Fig. S7D). By the PAP stage (∼24h), these proteins resolved into discrete puncta surrounding the nascent lumen, with PatJ/Pals1 facing the lumen and ZO-1 oriented away from it (Fig. 4J, Fig. S7E). Pearson’s correlation analysis showed again that PatJ colocalised more strongly with Pals1 than with ZO-1 (Fig. 4K). Collectively these data indicate that lumen initiation is temporally coordinated with the spatial reorganization of TJs and the VMZ.

### VACs transport apical membrane from the ECM-facing cortex to the AMIS

Next, we examined the spatio-temporal relationship between PatJ and apical cargo. As expected, at the inverted stage (16-18h), ezrin localised to the ECM-facing cortex while PatJ formed a double-puncta pattern at the periphery of the cell-cell contact interface (Fig. 5A). By ∼20h, ezrin was largely depleted from the ECM-facing cortex and instead accumulated in large (1-2 µm) intracellular vacuoles reminiscent of VACs (Fig. 5B). These vacuoles appeared to dock and fuse at the AMIS, accompanied by separation of PatJ into two discrete puncta, consistent with the onset of lumen initiation and junction assembly (Fig. 5C). By 24h, most cell doublets exhibited a PAP/nascent lumen phenotype, with ezrin labeling the luminal domain and PatJ puncta surrounding it (Fig. 5D). Together, these observations indicate that PatJ becomes enriched at the AMIS immediately prior to the arrival of VAC-like organelles carrying apical cargo.

**Figure 5:**
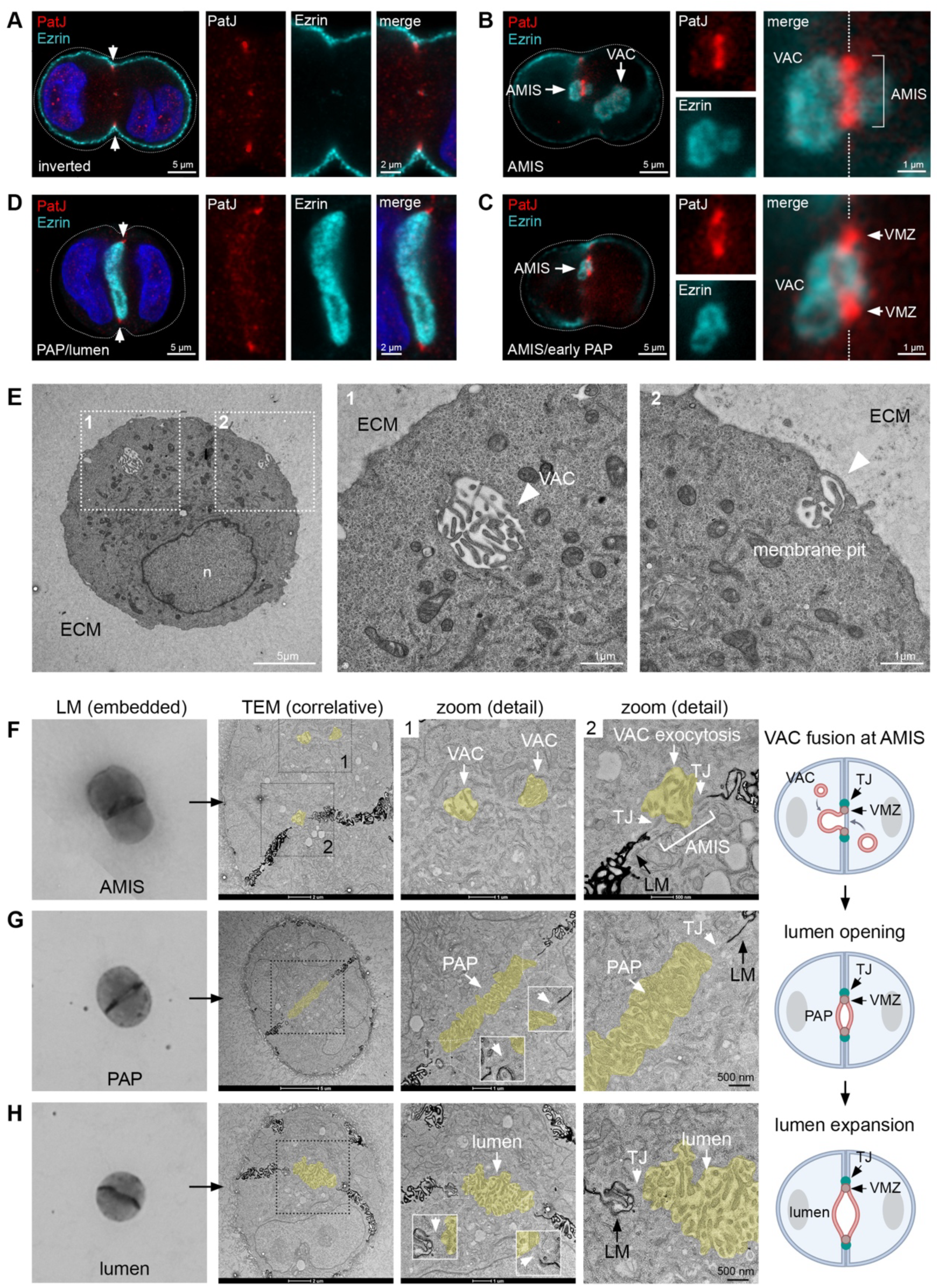
**VACs fuse at the AMIS to initiate lumen assembly** (A-D) Representative Airyscan images of inverted (A), AMIS (B), early PAP (C), and PAP/open lumen (D) stages. Cells were stained with anti-PatJ and anti-ezrin antibodies. Intracellular VACs (B) and VAC fusion events with the AMIS (B and C) are shown. (E) TEM micrograph of an 18h cell doublet showing an intracellular VAC-like organelle and a microvilli-containing membrane pit at the ECM-facing plasma membrane. (F-H) MDCK cells embedded in Matrigel for 24h were processed for correlative light and serial section TEM. Transmitted light images (left column) and representative correlative TEM micrographs of AMIS (F), PAP (G), and open lumen (H) stages are shown. The EM contrast of the lateral membrane (LM) was enhanced by osmium/tannic acid impregnation. Intracellular VACs and a VAC fusion event at the AMIS are shown (F). The VAC fusion site and the luminal domain is surrounded by TJs, which appear electron lucent (F-H). See also Figure S9.

To validate the identity of these VAC-like organelles, we analysed inverted stages by TEM. Large vacuoles containing apical microvilli were readily detected in early-stage cysts and closely resembled VACs observed in calcium switch assays (Fig. 5E). In addition, membrane pits containing microvilli-like protrusions were detected at the ECM-facing plasma membrane, potentially representing sites of apical membrane internalization (see also Fig. 6). To support these findings and better understand how VACs fuse with the AMIS, MDCK cells embedded in Matrigel for 24h were processed for correlative light and electron microscopy (CLEM). Cells were aldehyde-fixed and stained with osmium tetroxide/tannic acid, which enhanced contrast at the lateral but not the apical membrane and rendered apical cell junctions electron-lucent (Fig. S9A). This allowed us to unambiguously detect sites of lumen initiation and junction assembly without having to rely on genetically encoded EM probes. Cell doublets in the inverted, AMIS, PAP and early lumen stages were identified by transmitted light microscopy and subsequently imaged using serial section TEM (Fig. 5F-5H, Fig. S9B). As expected, multiple VACs as well as VAC fusion events at the AMIS were observed (Fig. 5F, S9B-S9D). VAC fusion sites were confined by cell junctions, which appeared as electron-lucent, flattened membranes distinct from the electron-dense, interdigitated lateral membranes. At PAP and early lumen stages, the lateral membrane remained strongly stained, whereas the microvilli-rich apical surface was unlabeled (Fig. 5G, 5H). Cell junctions consistently demarcated the boundary between the lumen and the lateral membrane. Taken together these data provide ultrastructural evidence that VAC exocytosis at the AMIS drives lumen initiation and is temporally coordinated with the assembly of apical cell junctions.

**Figure 6:**
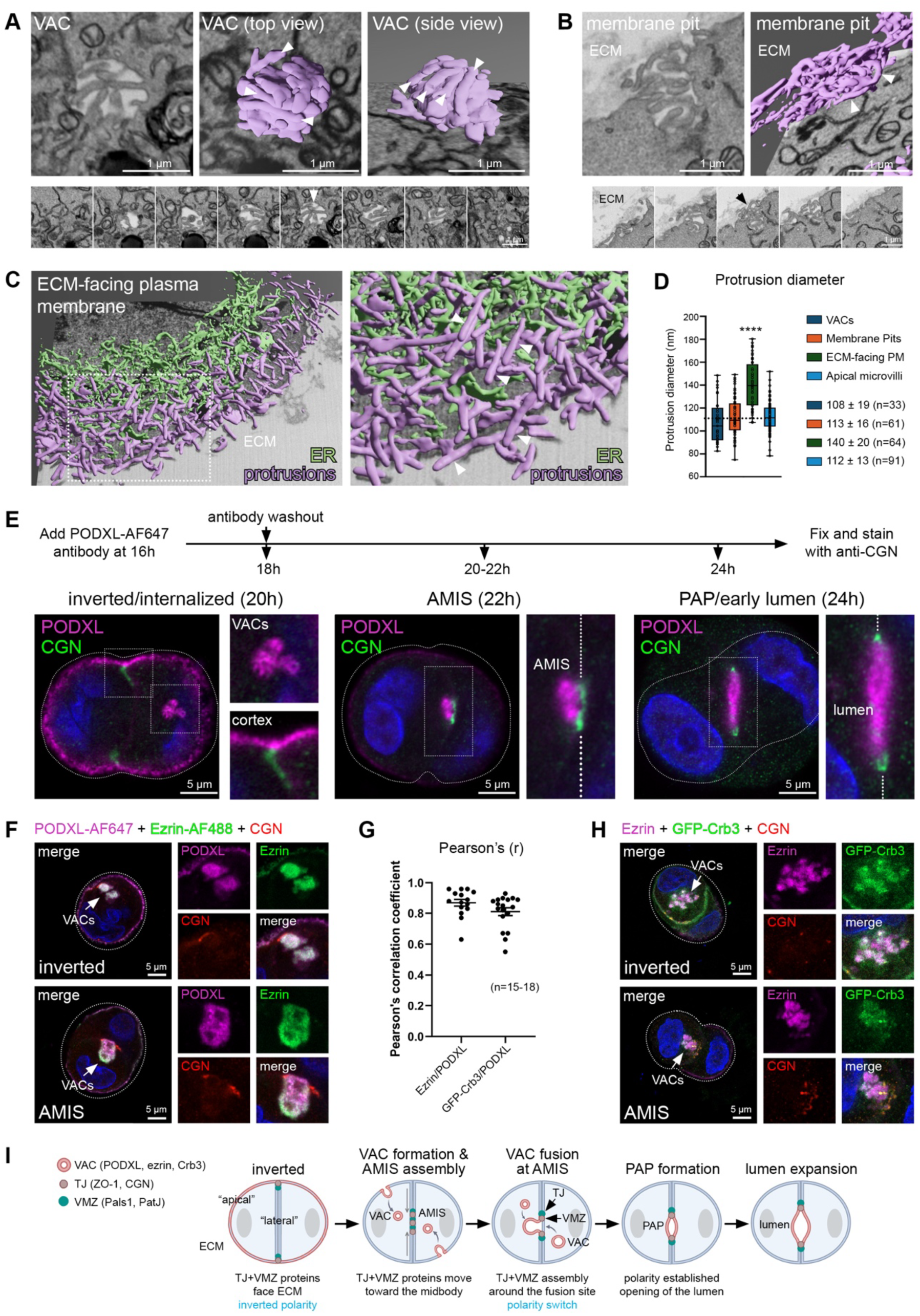
**VACs are transcytotic organelles that deliver a preassembled apical cortex to the AMIS** (A) FIB-SEM and 3D segmentation of an intracellular VAC. Microvilli are indicated by arrowheads. See also Figure S10 and Movies S1 and S2. (B) FIB-SEM and 3D segmentation of a microvilli-rich membrane pit at the ECM-facing plasma membrane. Microvilli are indicated by arrowheads. See also Figure S10 and Movie S3. (C) FIB-SEM and 3D segmentation of the ECM-facing plasma membrane. Membrane protrusions (white arrowheads) and ER membranes are shown. See Movie S4. (D) Quantification of protrusion diameters observed in VACs and membrane pits compared to protrusions at the bulk ECM-facing cortex and apical microvilli observed in mature MDCK monolayers. **** p<0.0001, Student’s t-test (n=33-91). (E) Transcytosis of PODXL was analysed using an antibody uptake assay. AF647-labelled PODXL antibody was added at 16h and washed out at 18h. Cells were fixed between 18h and 24h and stained with antibodies against CGN. Representative inverted, AMIS, and PAP stages are shown. (F) Transcytosis of AF647-labelled PODXL was analysed using an antibody uptake assay as in (E). Cells were fixed between 18h and 24h and stained with antibodies against CGN and a pre-labeled Ezrin-AF488 antibody. Inverted and AMIS stages are shown. See also Figure S11A and S11B. (G) Pearson’s correlation coefficients of ezrin/PODXL and GFP-Crb3/PODXL colocalisation in early-stage cysts based on data shown in Figure 6F and Figure S11C. n = 15-18 cysts each. (H) Inverted and AMIS stages of MDCK cells stably transfected with GFP-Crb3 stained with anti-ezrin and anti-CGN antibodies. See also Figure S11D. (I) Proposed new model of *de novo* lumen formation in MDCK cells embedded in Matrigel.

To visualize the 3D ultrastructure of VACs and explore how and where they are formed, we imaged PAP/early lumen stages using focused ion beam-SEM (FIB-SEM) at a resolution of 10×10×30 nm (Fig. 6A-6C, Fig. S10). Within the reconstructed volume, we identified multiple small, discontinuous “mini-lumens” and two VAC-like organelles (Fig. S10A, S10B). Three-dimensional segmentation revealed that these vacuoles contained densely packed arrays of protrusions characteristic of apical microvilli (Fig. 6A, Fig. S10B, Movies S1 and S2). Strikingly, several micrometer-scale membrane invaginations enriched in similar protrusions were observed on the ECM-facing plasma membrane (Fig. 6B, Fig. S10C, Movie S3), whereas the surrounding cortex outside these pits displayed sparser and morphologically distinct protrusions (Fig. 6C, Movie S4). Quantitative analysis showed that protrusions within VACs and membrane pits had an average diameter of ∼110 nm, closely matching apical microvilli in mature MDCK monolayers and previous reports (Fig. 6D) ^54,55^. By contrast, protrusions emanating from the ECM-facing plasma membrane were larger (∼140 nm), suggesting that apical microvilli are enriched in membrane pits but largely absent from the surrounding cortex. Together, these data demonstrate that VACs contain bona fide apical microvilli and likely originate from ECM-facing membrane pits that concentrate and internalize a preassembled apical cortex.

To directly test whether VACs deliver apical membrane from the ECM-facing cortex to the AMIS, we analysed PODXL trafficking using an antibody uptake assay. Fluorescently labeled PODXL antibody was added to MDCK cells at 16h in Matrigel and washed out at 18h. Cells were fixed at 18-24h and costained with anti-CGN antibody to visualize TJs (Fig. 6E). At 18h PODXL labeling was confined to the ECM-facing plasma membrane (not shown). At ∼20h PODXL started to accumulate in large intracellular vacuoles, which at ∼22h appeared to dock at the AMIS marked by CGN. At 24h PODXL was exclusively associated with the lumen surrounded by TJs, indicating transcytosis of PODXL from the ECM-facing plasma membrane to the AMIS (Fig. 6E). Moreover, ezrin, which interacts directly with PODXL ^56,57^, and GFP-Crb3 efficiently colocalized with internalized PODXL antibody in VAC-like organelles, at the AMIS, and at nascent lumens (Fig. 6F, 6G, Fig. S11A-S11C). GFP-Crb3 also colocalized with ezrin during transit and at the AMIS (Fig. 6H, Fig. S11D). We concluded that apical proteins are co-internalized from the ECM-facing cortex into VAC-like organelles that are subsequently transported to the AMIS (Fig. 6I).

### PatJ is required for VAC exocytosis at the AMIS and lumen initiation

To investigate the function of PatJ in lumen morphogenesis, we generated PATJ CRISPR/Cas9 MDCK knockout (KO) cells (Fig. S12A-S12D). Loss of PatJ expression was confirmed by Western blotting and immunofluorescence, which also revealed a marked reduction in both junctional and total Pals1 levels (Fig. 7A-7C). PatJ loss also significantly decreased ZO-1 levels at bicellular junctions and F-actin intensity at the apical domain (Fig. S12E-S12H). This supports earlier work demonstrating a function for PatJ and Pals1 in TJ formation ^38,39,42,44^ and further suggests that PatJ and Pals1 regulate the organisation of the apical cortex. This prompted us to examine the apical surface of MDCK monolayers directly by scanning EM (Fig. 7D). Wild-type cells displayed a dome-shaped apical surface densely covered with microvilli. By contrast, the apical surface of PATJ KO cells appeared flatter and less convex, and microvilli were sparser and more evenly distributed. Interestingly, wild-type cells exhibited radial arrays of elongated microvilli in the peripheral marginal zone that formed intercellular bundles with microvilli of neighboring cells (Fig. 7D, Fig. S13A, S13B). Automated segmentation revealed that both the density and the orientational alignment of these “marginal zone rods” were significantly reduced in PATJ KO cells (Fig. 7E, 7F, Fig. S13C-S13F). Moreover, rod widths were consistent with the dimensions of microvilli and were unchanged in the absence of PatJ (Fig. S13G). These findings indicate that PatJ regulates the architecture of the marginal zone and promotes the formation of ordered radial arrays of microvillar bundles across apical cell-cell contacts (Fig. 7G).

**Figure 7:**
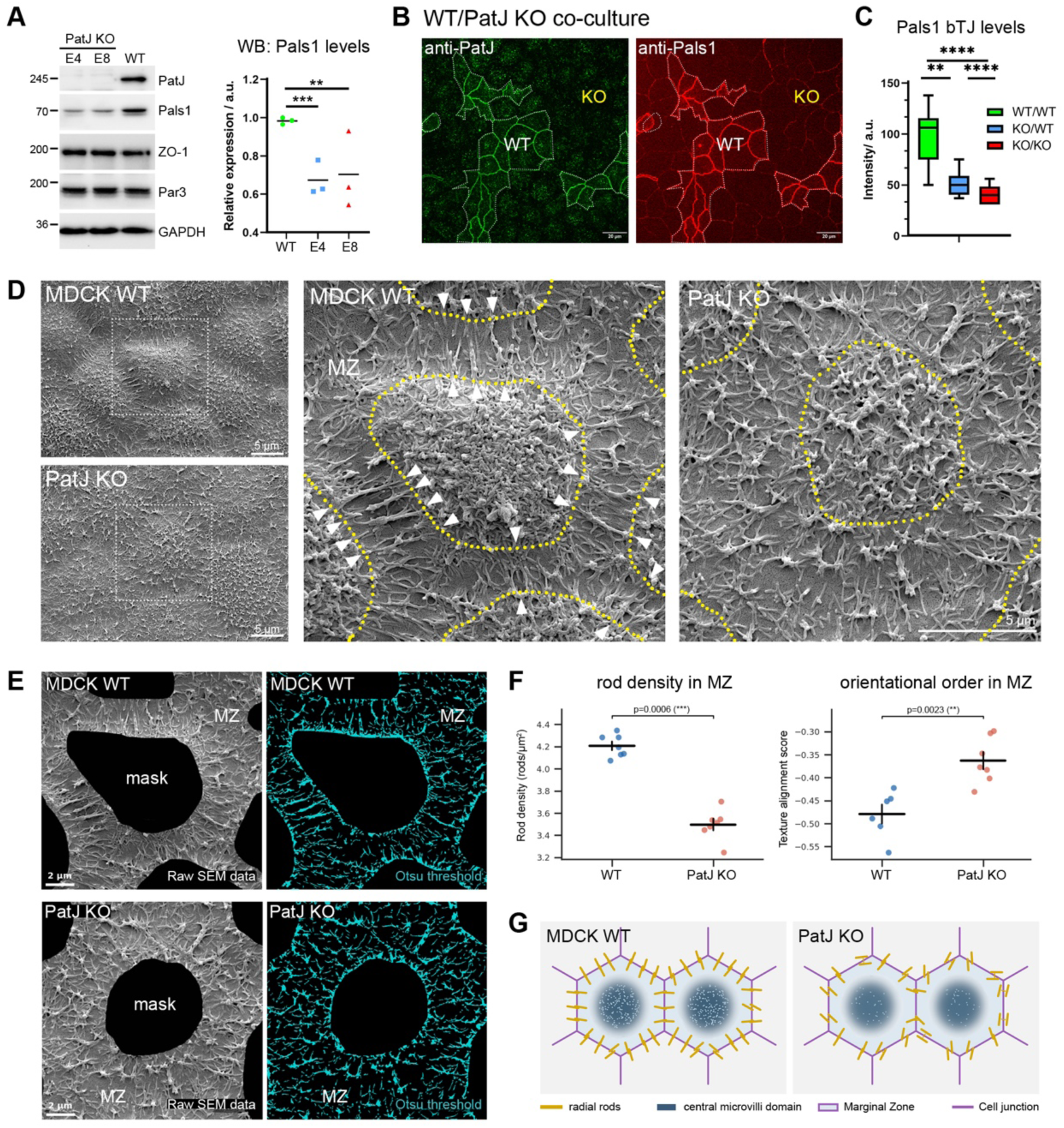
**PatJ controls the organisation of the marginal zone** (A) Western blot analysis of PATJ KO cells (clones E4 and E8) using the indicated antibodies. PatJ was probed with an antibody against the PDZ4 domain. Pals1 expression was quantified by densitometry. ** p<0.01, *** p<0.001, Student’s t-test (n=3). See also Figure S12. (B) MDCK WT and PATJ KO cells were co-cultured in a 1:1 ratio on Transwell filters for 12 days. Cells were fixed and stained with anti-PatJ and anti-Pals1 antibodies. Maximum intensity projections of confocal image stacks are shown. (C) Quantification of junctional Pals1 intensity based on data shown in (B). ** p<0.01, **** p<0.0001, Student’s t-test (n=20 junctions/condition). (D) MDCK WT and PATJ KO cells were cultured on Transwell filters for 16 days and processed for SEM. Representative micrographs are shown. Note the presence of rod-shaped cross-bridges (white arrowheads) in the marginal zone (MZ; bounded by yellow dotted lines) of WT cells. See also Figure S13A and S13B. (E) Automated thresholding and segmentation of rod-shaped microvilli in the MZ. The central microvilli-rich domain was excluded from the analysis (indicated as a black mask). See also Figure S13C. (F) Quantification of rod density and texture alignment in the MZ. Rod density was quantified based on a two-stage pipeline involving thresholding and segmentation. Texture alignment was performed on raw SEM images and measures the orientational coherence of all objects in the MZ; a negative score indicates high orientational order, while a score nearer to zero indicates a more random orientation. Error bars represent mean ± SEM. Statistical analysis was performed using a two-sided Mann-Whitney U test. *** p<0.001; **p<0.01 (n=6-7 micrographs/condition; 30-40 cells/condition). See also Figure S13D-S13F. (G) Model depicting the function of PatJ in regulating the density and radial organization of microvilli-based rods in the marginal zone.

Next, we addressed the function of PatJ, and hence the marginal zone, in lumen formation. Loss of PatJ resulted in a severe and fully penetrant multi-lumen phenotype, with nearly all KO cysts forming multiple lumens instead of a single central lumen (Fig. 8A, 8B). PALS1 KO cells exhibited a similar phenotype, consistent with previous work ^39,44^. To determine whether this defect arises from defective lumen initiation, we analysed the trafficking of apical proteins in early-stage cysts using Airyscan and 3D Structured Illumination Microscopy (Fig. 8C-8E). In WT cells ezrin and internalized PODXL antibody localized to the luminal domain surrounded by TJs labeled with CGN. By contrast, in PATJ KO cells apical proteins accumulated in large intracellular vacuoles and failed to be delivered to the center of the cyst. Loss of PatJ also affected the localization of CGN, which appeared diffuse or as scattered puncta in the cytoplasm. To quantify the lumen initiation defect, we measured the displacement (offset index) of PODXL-positive vacuoles relative to the center of the cell doublet (Fig. 8F, 8G). PATJ KO cells showed a significantly higher offset index than WT cells, indicating that PatJ is required for the incorporation of VACs into the AMIS, and thus for lumen initiation.

**Figure 8:**
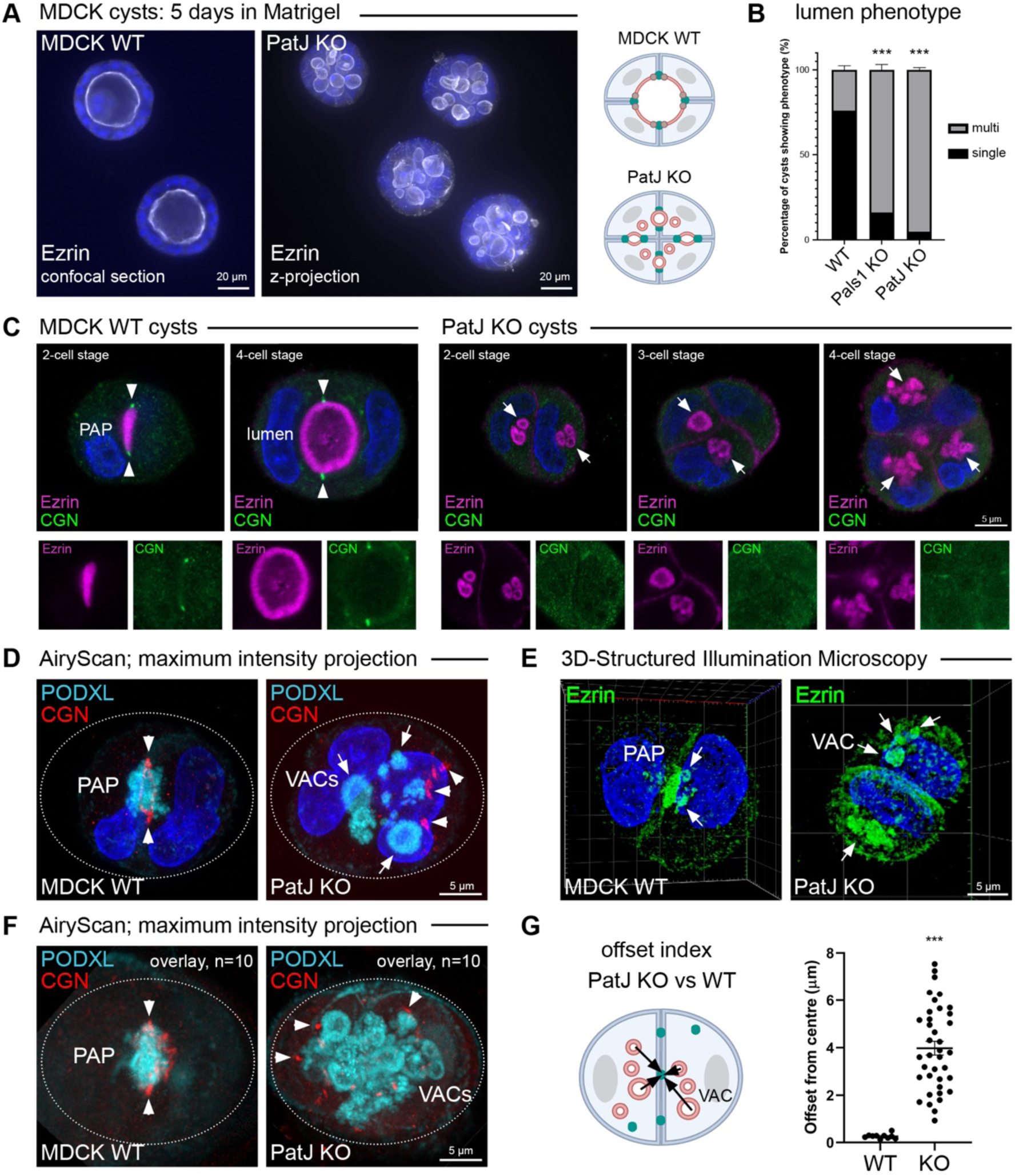
**PatJ is required for lumen initiation** (A) Representative confocal images of MDCK WT and PATJ KO cysts embedded in Matrigel for 5 days. Cysts were stained with anti-ezrin antibody. The PATJ KO cysts are shown as maximum intensity z-projections. (B) Quantification of the data shown in (A). Cyst phenotypes (single or multi-lumen) were quantified from three independent experiments. Error bars represent mean ± SEM. ***p ≤ 0.001, Student’s t-test (n=100 cysts each). (C) Representative Airyscan images of WT and PATJ KO cysts cultured in Matrigel for 24h or 48h. Cysts were stained with anti-ezrin and anti-CGN antibodies. 2-cell, 3-cell, and 4-cell stages are shown. White arrowheads indicate TJs labeled by CGN. White arrows indicate VACs in the cytoplasm. (D) Representative Airyscan images of WT and PATJ KO cysts cultured in Matrigel for 24h. Fluorescently labeled anti-PODXL antibody was added to the culture at 16h and washed out at 18h. Cysts were co-stained with anti-CGN antibodies. Note the accumulation of PODXL in intracellular VACs (white arrows) and the presence of CGN puncta in the cytoplasm (white arrowheads) in PATJ KO cells. (E) Representative 3D SIM reconstructions of WT MDCK and PATJ KO cysts cultured in Matrigel for 24h. Cells were stained with anti-ezrin antibodies. Note the accumulation of ezrin in intracellular VACs (white arrows) in PATJ KO cells. (F) 10 WT and 10 PATJ KO PAP stages (24h) stained with anti-PODXL and anti-CGN antibodies were overlayed and aligned. (G) Quantification of the “offset index”, defined as the distance of PODXL/ezrin-stained structures from the center of the cell doublet. Error bars represent mean ± SEM. ***p ≤ 0.001, Student’s t-test (n=10 cysts each).

It has been shown that PatJ connects the TJ to the apical cortex and thereby stabilizes the apical-lateral border ^42,58^. To test if the loss of this physical linkage causes the lumen initiation defect, we generated PATJ KO cells stably re-expressing various forms of mCherry-tagged PatJ (Fig. 9A, Fig. S14A). We reasoned that the L27 domain, which binds Pals1, and the PDZ6 domain, which binds ZO proteins, would be minimally required to restore PatJ function ^42,58^. Unexpectedly, none of the PatJ constructs — including full-length PatJ — rescued the multi-lumen phenotype of PATJ KO cells (Fig. 9B, Fig. S14B). Moreover, ectopically expressed PatJ failed to localise to apical cell junctions and did not restore junctional Pals1 levels (Fig. S14C–S14E). This suggests that re-expression of PatJ is insufficient to reconstitute the Crumbs complex in PATJ KO cells, likely due to markedly reduced ZO-1 and Pals1 levels at apical junctions (Fig. 7A-C, Fig. S12E). To connect the TJ directly to Pals1 and bypass the functions of PatJ’s PDZ domains, we introduced into PATJ KO cells a chimeric construct in which the L27 domain of PatJ was fused to ZO-1 (Fig. 9A, Fig. S14A) ^42^. Interestingly, the chimeric PatJ-ZO1 construct was efficiently recruited to TJs, restored junctional Pals1 levels, and fully rescued the multi-lumen phenotype of PATJ KO cells (Fig. 9B, Fig. S14B–S14E). Furthermore, the chimera co-localised with CGN at the AMIS and restored the trafficking of apical proteins to the site of lumen initiation (Fig. 9C). This shows that direct tethering of the L27 domain to the ZO scaffold is sufficient to restore Pals1 localisation and marginal zone function. Taken together, the data indicate that PatJ links the TJ to the apical cortex and thereby functions as a key organizer of the apical-lateral border. In its absence, the apical marginal zone is disrupted and the TJ belt becomes destabilized, preventing VAC exocytosis at the AMIS and disrupting lumen initiation (Fig. 9D).

**Figure 9:**
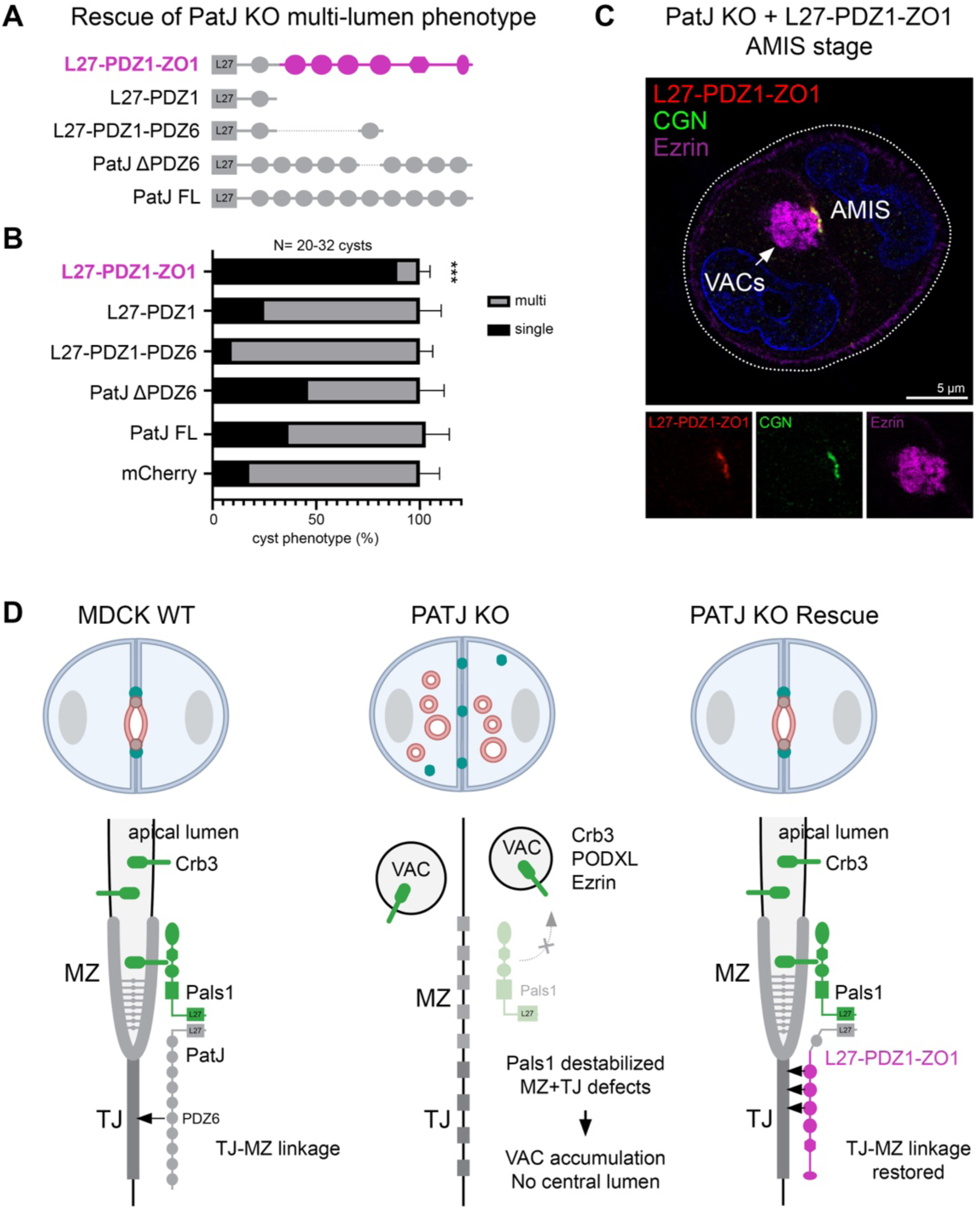
**PatJ connects the TJ to the apical cortex to initiate lumen formation** (A) PATJ KO cells were stably transfected with mCherry (control), the indicated PatJ constructs, or a chimeric PatJ L27-PDZ1-ZO1 construct. All constructs were expressed as N-terminal mCherry fusion proteins. See also Figure S14A. (B) Quantification of lumen formation in PATJ KO rescue cell lines. Single or multi-lumen phenotypes were quantified from three independent experiments. Error bars represent mean ± SEM. ***p ≤ 0.001, Student’s t-test (n=20-32 cysts each). See also Figure S14B. (C) Representative AMIS stage of PATJ KO cells stably transfected with the chimeric PatJ L27-PDZ1-ZO1 construct, fixed and stained with anti-PODXL and anti-CGN antibodies. (D) Model illustrating the function of PatJ and the marginal zone in lumen initiation. MZ=marginal zone; TJ=tight junction.

## Discussion

We show here that lumen initiation is driven by large micron-sized organelles, termed vacuolar apical compartments (VACs), which contain a preassembled microvilli-rich cortex and fuse at the AMIS to create a nascent lumen. VAC fusion was temporally coordinated with the assembly of apical cell junctions and was dependent upon the Crumbs complex scaffolding protein PatJ. Our findings redefine the mechanisms of *de novo* lumen formation in MDCK 3D cultures and suggest broader implications for lumenogenesis in other epithelial systems.

Our imaging and proteomics analyses revealed that VAC fusion and cell-cell junction assembly are orchestrated by a highly dynamic and hierarchical protein interaction network (Fig. 3). Integral TJ membrane proteins, including JAM-A (F11R), claudin-1, and claudin-4, as well as all major Rho GTPases (e.g. Cdc42, Rac1, RhoA) became enriched with Pals1 at early time points (∼0.5-2h), presumably demarcating the arrival of VACs at the lateral membrane. By contrast, VMZ and TJ scaffolding proteins such as ZO-1, ZO-3, CGN, and Par3 were recruited later (∼2-4h), coinciding with the period when VAC fusion sites and lateral lumens were most prominent. Apical proteins became enriched at the final stages of polarity development (∼4-6h), when a continuous TJ belt was evident and most lateral lumens had repositioned to the apical cortex. These temporal recruitment profiles are consistent with the established roles of Rho GTPases in junction assembly and maturation ^59–62^ and with established molecular interactions, including the role of claudins as ZO receptors and the ability of JAM-A to recruit ZO-1 and Par3 during TJ assembly ^63,64^. They are also consistent with and complementary to a previously reported time-resolved proteome of APEX2-tagged ZO-1 and with data showing that PatJ is required for the coalescence and condensation of ZO-1 along the apical-lateral interface ^42,65^. Together, these findings support a model of junctional maturation in which TJ and VMZ proteins become progressively concentrated and confined at the apical-lateral border, ultimately positioning the VMZ between the TJ and the apical cortex.

We show here that VACs function as apical precursor organelles for lumen initiation in MDCK 3D cultures (Fig. 6I). VACs were originally identified in calcium-depleted MDCK cells ^47,48^, but similar structures have since been observed in multiple cell types under physiological conditions ^66–68^, demonstrating that they are not artifacts of calcium depletion. More recently, VAC-like organelles have also been reported in several developmental systems. For instance, isolated human embryonic stem cells form a perinuclear compartment termed the “apicosome” ^17,49^. Apicosomes contain microvilli and apical proteins such as PODXL, are enriched in calcium, and mediate lumen initiation and expansion during mouse blastocyst development and in human stem cell-derived epiblast models ^12,13,69^. Large intracellular organelles containing apical markers have also been described during lumen formation in the mouse axial mesoderm ^70^. Moreover, intracellular VAC-like structures accumulate upon loss of Cdc42 in the mouse pancreas ^71^ and in microvillus inclusion disease, which is caused by mutations in the apical motor protein MYO5B ^72^. Interestingly, in the *Drosophila* midgut, pre-enterocytes establish an apical pole well before they are incorporated into the intestinal epithelium ^73,74^. This preassembled apical compartment (PAC) is an apical cavity surrounded by cell-cell junctions with neighboring enterocytes and shares key features with the apical luminal surface formed in the calcium switch assay. Together with our findings, these observations suggest that VAC-like compartments contribute to apical membrane assembly in diverse cellular and developmental contexts.

Current models of *de novo* lumen formation in MDCK cells propose that apical cargo is delivered to the AMIS via vesicular transport (Fig. 4A). Such carriers contain apical Rab proteins including Rab3, Rab8 and Rab11 and are proposed to be of recycling endosomal nature ^2–4,16^. However, our data and careful examination of the MDCK cell literature indicates that PODXL-containing transport carriers do not always appear vesicular. In fact, Crb3 ^40^, Rab11 ^19^, IRSp53 ^25^, Rab35 ^24^, and the synaptotagmin-like protein Slp2 ^22^ (to name a few) often colocalise with PODXL in large intracellular vacuoles highly reminiscent of VACs. In addition, apical cargo frequently accumulates in such vacuoles upon perturbation of the apical trafficking machinery, TJ proteins, or polarity factors ^19,21,22,24,25,40,75,76^. This indicates that VACs have been observed previously but were not recognized as a distinct organelle class. Because VACs can only be reliably resolved by TEM or high-resolution light microscopy, their prevalence may have been underestimated. Indeed, only two studies have examined the ultrastructure of early-stage MDCK cysts ^25,77^, and these analyses sampled relatively small cellular volumes, possibly explaining why VAC-like structures were not reported before.

Altogether, we propose that VACs act as transcytotic trafficking devices that deliver a preassembled apical cortex to the AMIS, enabling rapid and efficient lumen initiation (Fig. 6I). Our data suggest that VACs arise through the internalization of microvilli-enriched membrane pits from the ECM-facing plasma membrane. How apical proteins become concentrated in these pits and subsequently internalized remains unclear. Similarly, the molecular mechanisms underlying VAC trafficking and fusion at the AMIS remain to be determined. Based on the literature and our proteomics data, we considered whether apical Rab proteins and the exocyst complex might contribute to VAC trafficking and fusion ^4^ (Fig. S15). Consistent with a role for the exocyst in VAC exocytosis, pharmacological inhibition of EXO70 using endosidin-2 altered lumen morphology and produced a mild multi-lumen phenotype. By contrast, CRISPR-mediated inactivation of Rab3, Rab11, Rab25, or Rab35 did not noticeably perturb lumenogenesis, indicating substantial redundancy within the Rab family in apical membrane trafficking, at least in MDCK cells ^78^. We also assessed the role of Rho/actomyosin signaling in lumen initiation. Whilst lumen shapes were altered, neither activation nor inhibition of Rho or myosin-II activity impaired lumen formation (Fig. S15) or rescued the lumen defects in PATJ KO cells (data not shown), supporting earlier work showing that junctional tension regulates lumen morphology but not assembly ^27,77,79^. Future studies should explore how VACs are transported to and fuse at the AMIS and address their functional significance in lumen forming tissues.

We demonstrate that VACs deliver Crb3, PODXL and ezrin to the AMIS. This agrees with previous data showing that Crb3 and PODXL are transported to the AMIS in a common transport carrier ^24,40^, but contrasts with findings that ezrin and PODXL are transported separately ^76^. Importantly, whilst endogenous Pals1 and PatJ were associated with VACs in the calcium switch assay, they did not localize to VACs formed in 3D cysts. Instead, Pals1 and PatJ accumulated at the AMIS before apical cargo including Crb3a arrived, suggesting that the full Crumbs complex is assembled only after VACs fuse at the AMIS. This differs from previous studies reporting that GFP-Crb3, Pals1 and PatJ are transported together to the AMIS ^40^. We find that even subtle overexpression of Crb3 promotes the recruitment of Pals1 onto the VAC membrane (data not shown), providing a possible explanation for the discrepancies between our findings and earlier reports.

Our data further indicate that AMIS formation requires TJ proteins, Pals1 and PatJ to translocate from peripheral, ECM-facing cell junctions to the center of the cell doublet. The arrival of VACs at the AMIS then triggers the formation of a junctional belt around the fusion site, with the VMZ facing the nascent lumen and the TJ facing toward the newly established lateral interface. These observations raise several important questions, including how junctional proteins relocate to the AMIS, how E-cadherin-based adhesions are cleared along the lateral interface to permit AMIS assembly and VAC fusion, and how VAC exocytosis and junction assembly are coordinated at the AMIS. Consistent with an important role of the VMZ in this process, loss of PatJ perturbed the recruitment of TJ proteins to the site of AMIS assembly and prevented the incorporation of VACs into the AMIS, resulting in severe defects in lumen morphogenesis (Fig. 9D). Moreover, and intriguingly, PatJ loss profoundly altered the structure of the apical marginal zone. In wild-type cells we discovered radial arrays of microvilli bundles connecting the apical surfaces of neighboring cells. These structures differed morphologically from intermicrovillar adhesion complexes found in intestinal epithelia ^80,81^ and were markedly disorganized in PATJ KO cells. We propose, therefore, that the Crumbs complex organizes the marginal zone to create a physical link between the apical cortex and the TJ, and that the “marginal cross-bridges” are a structural manifestation of this linkage. By mechanically coupling neighboring apical cortices, these structures may stabilize the apical-lateral border and thereby promote TJ belt assembly and lumen initiation. Of note, a chimeric construct consisting of the PatJ L27 domain fused to ZO-1 was sufficient to rescue the lumen formation defects observed in PATJ KO cells, directly supporting the model that the physical tethering of the TJ to the apical cortex — rather than TJ assembly *per se* — is the critical determinant of lumen initiation (Fig. 9D). Consistent with this interpretation, although loss of TJ proteins, such as ZO-1 or Par3, often results in multi-lumen phenotypes ^75,82,83^, a fully functional TJ belt does not appear to be strictly required for single lumen formation in Matrigel cultures ^27,84^. In summary, our findings identify PatJ as a key structural and functional scaffold of the marginal zone, which promotes the assembly of a circumferential TJ belt and establishes the structural framework required for VAC-mediated lumen initiation.

## Acknowledgements

We thank Hi-Tech Instruments Pte Ltd and Hitachi High-Tech Corporation for the FIB-SEM analysis. We are grateful to Thomas Weide for providing Pals1 KO cells, Alf Honigmann for providing mNeonGreen-ZO1 cells and mCherry-tagged PatJ constructs, Andre Le Bivic for sharing the PatJ L27 antibody, Arnaud Echard for the GFP-Crb3 construct, and Sandra Citi for providing anti-cingulin and anti-ZO1 antibodies. We thank Effie Iliffe-Moon for help in designing figures and members of the NTU Institute of Structural Biology (NISB) for technical support. This work was supported by a Singapore Ministry of Education (MOE) Academic Research grant Tier2 (MOE-T2EP30121-0019) to A.L. E.M. and R.G. were supported by an FNR AFR Bilateral Singapore grant (11823257) to G.D. and A.L.

## Author contributions

S.G. performed all MDCK cyst experiments, E.M. performed and analysed the proteomics work, generated Rab overexpression cell lines, and carried out TEM analysis of Rabs, Y.C. generated and characterized PATJ KO and rescue cell lines and performed calcium switch experiments, N.A. performed colocalization analysis in 2D cultures, S.S. segmented the FIB-SEM data, K.E.L. performed SEM and prepared samples for FIB-SEM, R.G. acquired MS data, B.H. performed TEM analysis, G.D. supervised E.M. and R.G. during the early stages of the project and overlooked the proteomics analysis, A.L. supervised S.G., E.M., Y.C., N.A., R.G., and B.H., conceived the project, performed TEM and CLEM experiments, and wrote the manuscript.

## Code Availability

All Fiji/ImageJ Macros can be found here: https://github.com/Microscopy-Cluster-NUS/Microvilli_ER_vEM

## Materials and Methods

### DNA constructs, antibodies and cell lines

The Pals1-APEX2-EGFP and Pals1-GFP MDCK-II cell lines have been described previously ^43,44^. C1-EGFP-hPMCA2w/b (plasmid #47586), C1-EGFP-Rab3a (plasmid #49542, GenBank: AF498931) and C1-EGFP-Rab35 (plasmid #49552, GenBank: NM_006861) were purchased from Addgene. C1-EGFP-Rab1a (GenBank: NM_00416) and C1-EGFP-Rab6a (GenBank: NM_198896) were provided by Lu Lei. C1-EGFP-Rab11a (GenBank: NM_001003276) was provided by Ben Nichols. The cDNA for Rab25 (GenBank: BC033322, IMAGE clone: 4775725) was provided by the Protein Production Platform, Nanyang Technological University. To produce APEX2-EGFP versions of Rab1a, Rab3a, Rab6a, Rab11a, Rab25 and Rab35, the cDNAs were cloned into the C1-A2E vector ^43^ using FastCloning ^85^. The pX459 CRISPR/Cas9 vector (Addgene plasmid #62988) was provided by Xavier Francesc Roca Castella. Single guide RNA sequences were inserted using the *BbsI* site. All constructs were verified by Sanger sequencing. Rab3 (Rab3a/b/c/d quadruple KOs), Rab11 (Rab11a/b double KOs), Rab25, and Rab35 MDCK-II KO cells were purchased from RIKEN BRC^78^. MDCK-II cells expressing endogenously tagged mNeonGreen-ZO1 and mCherry-tagged PatJ constructs were a kind gift from Alf Honigmann and were described previously ^42^. The PatJ anti-L27 antibody was kindly provided by Andre Le Bivic. Rabbit anti-cingulin and rat anti-ZO-1 antibodies were a kind gift from Sandra Citi. GFP-Crb3 was kindly provided by Arnaud Echard.

### Cell culture

MDCK-II cells were cultured in high glucose DMEM with pyruvate and GlutaMAX^TM^ supplement (Gibco, 10569010) supplemented with 10 % FBS, 100 units/mL penicillin, and 100 μg/mL streptomycin at 37 °C and 5 % CO_2_. Cells were allowed to grow to confluence before passaging. Trypsinisation was performed by washing with Phosphate Buffered Saline (PBS) pH 7.4 (Gibco) followed by incubation with 0.25% trypsin-EDTA (Gibco) until detached. For monolayers requiring complete polarisation, MDCK-II cells were seeded into 0.4 µm-pore Transwell® filter inserts (Corning) at an initial density of ∼2 x 10^5^ cells/cm^2^. Media was changed every two days by aspirating out old media and adding in fresh media. Cells were grown on the filter inserts for 10-14 days to achieve full polarisation.

### Generation of stable cell lines and CRISPR/Cas9 knockout lines

Cells were transfected with Lipofectamine 3000 (Invitrogen) according to the manufacturer’s instructions. For the generation of stable cell lines expressing APEX2-EGFP fusion constructs, MDCK-II cells were selected with 500 µg/mL geneticin (GIBCO) for at least 14 days post-transfection and subsequently FACS sorted for GFP positive cells. Pals1 MDCK KO cells have been described previously ^44^. For the generation of CRISPR/Cas9 PatJ knockout cells, three sgRNAs (see Supplementary File S2) were designed from the Benchling App, each targeting specific 20 nt-long sequences in early exons (exon 4, 11, 16) of the canine *PatJ* gene. These gene locations correspond to the L27 (sgRNA2), PDZ3 (sgRNA1), and PDZ4 (sgRNA3) domains of PatJ, respectively. The sgRNAs were inserted into the pX459 backbone, which also encodes the tracrRNA scaffold and Cas9 protein, and transfected into MDCK-II cells. MDCK-II cells were co-transfected with C1-EGFP and two pX459 sgRNA plasmids in a 1:4 ratio. Cells were selected with 500 µg/mL geneticin (Gibco) for at least 14 days post-transfection. The cells were then GFP FACS sorted as single cells into 96 well plates. Single cell clones were expanded and screened by Western blotting. Three PATJ KO lines (E4, E6, E8) were selected from cells co-transfected with sgRNA1 and sgRNA3. Their genomic DNA was extracted to amplify the sgRNA1 (exon 11) target site by PCR. The purified PCR products were submitted for DNA sequencing. The three KO lines were genetically identical, having a 247 bp insertion in exon 11, resulting in a premature stop codon in the PDZ3 domain.

### MDCK 3D cultures

For 3D cyst assays, MDCK-II cells were split sparsely (∼1:10-1:50) to obtain sub-confluent cell cultures. Matrigel® (Corning; Cat# 354230) of the required quantity was left at 4°C to thaw overnight. All steps using Matrigel® were performed on ice unless otherwise stated. The following day, glass-bottomed 8-well chamber slides (IBIDI) were coated with ∼4 µL Matrigel® per well and left undisturbed in the 37°C incubator to polymerise for 30-45 mins. In the meantime, MDCK-II cells were trypsinised and counted. Single cells were resuspended in 200 µL/well of 2% Matrigel® and seeded at a density of 30,000-50,000 cells/well. The 2% Matrigel® media was replenished every two days. Cysts were cultured for 16-20h for the inverted polarity and AMIS stages, 20-24h for the PAP and early lumen stages, and up to 4 days for the mature end-stage of lumen formation. For certain experiments, MDCK cells cultured in Matrigel were cultured in the presence of Blebbistatin (50 µM), Y-27632 (20 µM), Rhosin (20 µM), CNO3 (1 µg/ml), or Endosidin-2 (50 µM).

### Calcium switch assay

Low calcium media was formulated using DMEM without calcium chloride (Gibco, 21068028) supplemented with 3 μM calcium chloride, 110 mg/L (1 mM) sodium pyruvate, 3.97 mM L-alanyl-glutamine (GlutaMAX^TM^), 100 units/mL penicillin, 100 μg/mL streptomycin, and 5 % dialysed FBS. Normal calcium media was prepared by supplementing low calcium media with 1.8 mM CaCl_2_. MDCK-II cells were seeded at 50% density and grown to confluence overnight in 24-well plates on glass coverslips. Cells were washed twice with PBS and subsequently incubated in low calcium media overnight (16-24h) for complete disassembly of cell junctions. Media was then replaced with normal calcium media and cells were fixed and analysed at defined time points post-calcium addition.

### Immunofluorescence microscopy

MDCK monolayers and MDCK cells grown in Matrigel were washed thrice with PBS pH 7.4 followed by fixation with 1% paraformaldehyde (PFA) in PBS pH 7.4. Monolayers were fixed for 20 min at room temperature, whilst 3D cysts were fixed for 30 min at room temperature. 3D cysts were permeabilised in PBS pH 7.4 containing 0.3% Triton X-100 for 30 min, while monolayers were permeabilised in PBS 0.1-0.5% Triton X-100 for 10-15 min and washed thrice again with PBS. Samples were blocked with 10% FBS in 0.2 µm-filtered PBS pH 7.4 for at least 2h at room temperature or overnight at 4°C, and then incubated with primary antibodies diluted in antibody incubation buffer (0.2 µm-filtered PBS pH 7.4, 0.1% BSA, 0.01% Tween-20) for 2-3h at room temperature or overnight at 4°C. Samples were washed three times for 5 min each in antibody incubation buffer and further incubated for 1h with fluorescently conjugated secondary antibody diluted in antibody incubation buffer at room temperature in the dark (see Supplementary File S2 for all antibodies used). Samples were again washed thrice for 10 min each in PBS and stained for 10 min with 20 ng/mL DAPI (Sigma) diluted in PBS. For cells grown on Transwell® filter inserts (Corning), filter inserts were cut out with a sharp razor blade post blocking. For staining and washes, filter pieces were laid face-up on a piece of parafilm in a humidified chamber and overlaid with ∼25 µl of staining/wash solution. Glass coverslips were laid face-down over ∼40 µl of staining solutions on parafilm in a humidified chamber and were laid face-up on parafilm for washes. Filter pieces were placed face-up on glass slides, overlaid with mounting medium and covered with #1.5 (12 mm) glass coverslips (VWR). Cells were stained with DAPI (1 ug/mL in PBS) prior to mounting in mounting medium (Vectashield, Vector Laboratories H-1000). Coverslips were sealed with transparent nail polish and stored at 4°C. Monolayers grown on glass bottom dishes or Transwell® filters were imaged using the CorrSight spinning disk microscope (ThermoScientific) equipped with an Orca R2 CCD camera (Hamamatsu) using a 40x oil objective (NA 1.3, Zeiss, pixel size 0.163 µm) or a 63x oil objective (NA 1.4, Plan Apochromat M27, Zeiss, pixel size 0.1 µm) and standard filter sets. Pearson’s correlation analysis was performed on images acquired on a Zeiss 980 LSM with Airyscan 2 operated in super-resolution mode using a 63x oil objective (Zeiss, NA 1.4). 3D Structured Illumination Microscopy (3D-SIM) was performed on a Zeiss Elyra 7 with Lattice SIM. Confocal images were processed in Zeiss Zen software and ImageJ/Fiji ^86,87^.

### Live cell imaging

MDCK-II cells stably expressing Pals1-A2E were grown on glass-bottom dishes (MatTec Corp., Ashland, MA) and imaged live in low calcium media with a CorrSight spinning disk microscope (Thermo Fisher Scientific). Imaging was carried out at 37 °C, 5 % CO_2_ and 90 % humidity in a closed atmosphere chamber (IBIDI). Confocal z-stacks were acquired using a 63x oil objective (Plan Apochromat M27, Zeiss, NA 1.4) using standard filter sets. Focus was maintained by a hardware autofocus system (Focus Clamp). The laser output power and exposure times were set to a minimum.

### PODXL antibody uptake assay and co-labeling

In some experiments, primary antibodies were labelled using the FlexAble 2.0 Coralite Plus Antibody Labelling Kit (Proteintech Cat. No. KFA521, KFA523) according to manufacturer’s instructions. In brief, 1 µL of FlexLinker was added per 0.5 µg of primary antibody, diluted in the supplied FlexBuffer, mixed gently and incubated for 10-15 minutes in the dark at room temperature prior to use. Labelled PODXL antibody was added to the media at 16h and incubated for 2h before washing off. The cysts were then fixed at 2h intervals between the 18 to 24h time points, permeabilised, blocked, stained with phalloidin or additional primary and secondary antibodies. For co-staining of PODXL and Ezrin (both raised in mice), the antibodies were individually labelled with different conjugated dyes followed by the addition of FlexQuencher reagent, as per manufacturer’s recommendation for multiplexing with antibodies from the same species.

### Transmission electron microscopy

MDCK-II cells grown on glass bottom dishes (MatTec Corp., Ashland, MA) or in Matrigel were fixed with 2% glutaraldehyde (EMS) in 0.1 M cacodylate buffer (CB) (sodium cacodylate pH 7.4, 2 mM CaCl_2_) at room temperature for 5 min followed by incubation on ice for 1-2h. APEX2 labeling was performed as previously described ^43,50,88^. Cells were post-fixed with 1% osmium tetroxide (OsO_4_) (EMS) in CB containing 1% potassium ferricyanide (K_3_[Fe(CN)_6_]) (EMS). For TEM of 3D cysts, lateral membrane contrast was enhanced through incubation with 1% low molecular weight tannic acid (C_14_H_10_O_9_)_n_, (EMS) in CB for 1h at room temperature. In some instances, samples were *en bloc* stained with 1% uranyl acetate (UA) (EMS) in ultrapure water overnight at 4°C. Samples were dehydrated in a cold graded ethanol series (20%, 50%, 70%, 90%, 100%) for 2-5 min each, followed by two washes in 100% anhydrous ethanol (EMS) for 3 min each. Samples were then infiltrated with a Durcupan resin:ethanol gradient (1:2, 1:1, 2:1) for 45 min each followed by infiltration in 100% resin overnight. Five to eight more resin exchanges were performed for 45 min each at 45°C. Resin-embedded samples were polymerised at 60°C for 2 days. Samples were sectioned into 70-80 nm thick sections and mounted onto formvar and carbon-coated slot grids (EMS). Transmission electron microscopy (TEM) was performed using a Tecnai T12 TEM (FEI) operated at 120 kV. Electron micrographs were captured using a 4k x 4k Eagle CCD camera (FEI) and processed in ImageJ/Fiji.

### Focused Ion Beam SEM

For FIB-SEM, cells embedded in Matrigel for 24h were chemically fixed and processed following the osmium-thiocarbohydrazide-osmium (OTO) method, along with *en bloc* UA and lead aspartate staining before being embedded in resin ^89^. The specimen was trimmed into blocks less than 1000 µm^3^ in volume, mounted on aluminium pins, sputter-coated with platinum (LEICA EM ACE200), and imaged on a Hitachi NX9000 FIB-SEM. FIB slicing was performed at 30 kV acceleration voltage using a probe current of 12 nA and a slicing pitch/step size of 30 nm. The block face was imaged at 2 kV acceleration voltage using the BSE detector and an exposure time of 100 sec. The recorded images had a voxel size of 10 nm (x) x 10 nm (y) x 30 nm (z).

### FIB-SEM image processing, 3D segmentation and quantitative analysis

For 3D analysis, the full image stack (629 sections) was rescaled in ImageJ/Fiji to obtain isotropic voxels. The original axial spacing of 30 nm was interpolated to 10 nm to match the lateral pixel size, resulting in voxel size of 10 x 10 x 10 nm. Images were then inverted and adjusted for brightness and contrast, followed by Gaussian filtering and rebinning to improve visibility of fine structures and to reduce data size for subsequent processing. Three regions of interest (ROIs) were segmented: an intracellular VAC containing densely packed microvilli (slices 158-212), a microvilli-enriched membrane pit (slices 1-90), and an area of the ECM-facing cortex without membrane pit (slices 419-530). Microvilli-like structures and endoplasmic reticulum were segmented using the Trainable Weka Segmentation plugin in Fiji ^90^, with separate classifiers trained for each ROI based on representative pixel features. The resulting probability maps were normalized, converted to 8-bit, and thresholded to generate binary masks. Segmented structures were converted to 3D surface meshes and exported in Wavefront (.obj) format. Meshes were lightly smoothened and simplified in MeshLab ^91^ before final rendering and animation in Blender v4.3.4 (Blender Development Team, 2024) to visualize microvilli-like protrusions.

### Scanning Electron Microscopy

MDCK-II cells grown on Transwell filters for 16 days were fixed with 2% glutaraldehyde (EMS) and 2% EM grade paraformaldehyde (EMS) in 0.1 M cacodylate buffer (CB) (sodium cacodylate pH 7.4, 2 mM CaCl_2_) at room temperature for 5 min followed by incubation at 4°C for 2h. The samples were post-fixed with 1% OsO_4_ for 1h, dehydrated in a graded ethanol series (25%, 50%, 75%, 90% and 100%), dried using a critical point drier (LEICA EM CPD300), sputter coated with gold (LEICA EM ACE200), and examined at 5 kV using a Scanning Electron Microscope (Quanta 650FEG, FEI).

### Quantitative analysis of the marginal zone

The density of rod-shaped protrusions and their orientational coherence were quantified on SEM micrographs recorded at 5000x magnification (pixel size 13.48 nm) using a custom Python pipeline implemented in scikit-image. Seven images per condition (WT and PATJ KO) were analysed. The central apical microvilli-rich domain was masked and excluded from the analysis. Pixels with intensities below 25 (on a 0–255 scale) were classified as background, and connected background regions smaller than 5,000 pixels were removed by morphological filtering. Background subtraction was performed by applying a Gaussian blur (σ = 25 pixels, corresponding to ∼337 nm) to the normalised image (0–1 range) and subtracting the result from the original, clipping negative values to zero. This suppresses low-frequency illumination variation while preserving fine apical surface structures. An Otsu threshold was then computed on the background-subtracted pixel intensities and applied to produce a binary object mask. A size filter of 30–10,000 pixels² (∼0.005 – 1.8 µm²) was applied to restrict the analysis to objects with physical dimensions consistent with microvilli and microvilli rods. Object density was calculated as the number of retained objects per area (in µm²) for each image.

Texture alignment of the rod array was quantified independently of segmentation using the gradient structure tensor. The image was pre-smoothed with a Gaussian (σ_noise = 1.5 px) and Sobel gradients were computed in x and y. The outer product of the gradient vector was then integrated with a second Gaussian (σ_smooth = 12 px, ∼162 nm) to yield the per-pixel structure tensor J, from which the local dominant gradient orientation θ and coherence c = ((λ₁ − λ₂)/(λ₁ + λ₂))² were derived, where λ₁ and λ₂ are the tensor eigenvalues. Coherence ranges from 0 (isotropic texture) to 1 (perfectly oriented texture) and was used as a per-pixel weight. For each pixel in the marginal zone (500–2,500 nm from the apical mask boundary), the alignment score was computed as −cos(2(θ − φ)), where φ is the angle toward the nearest boundary pixel; this gives −1 for a rod oriented radially (perpendicular to the boundary) and +1 for a rod oriented tangentially (parallel to the boundary). Two complementary scores were computed: a simple coherence-weighted mean over all marginal zone pixels (simple band), and a mean computed after first binning pixels into a 2D grid of distance × normalised arc-length position along the boundary (arc-map grid). The arc-map grid corrects for the geometric bias introduced by straight boundary segments contributing disproportionately more pixels than curved regions; both methods gave concordant results. Grubbs outliers (G = 2.044 > G_crit = 2.020, α = 0.05) were excluded from alignment statistics. Statistical comparisons were performed using a two-sided Mann–Whitney U test (n = 6 WT, n = 7 PatJ KO for alignment; n = 7 per condition for density). All analyses were performed in Python using NumPy, SciPy, pandas, and scikit-image. Figures were generated with Matplotlib.

### Proximity biotinylation

APEX2 proximity biotinylation was carried out as described previously ^43,51^. Briefly, MDCK-II cells stably expressing Pals1-A2E were incubated in media containing 2.5 mM biotin phenol (Iris Biotech) for 30 minutes at 37 °C. Cells were washed twice with PBS and the APEX reaction was initiated by incubating with 250 mM H_2_O_2_ for 1 minute at room temperature. The APEX2 reaction was quenched by washing three times with the quencher solution (PBS containing 5 mM (±)-6-hydroxy-2,5,7,8-tetramethylchromane-2 carboxylic acid (Trolox), 10 mM sodium ascorbate, 10 mM sodium azide) prior to lysis in ice-cold lysis buffer (1 % sodium deoxycholate in 50 mM ammonium bicarbonate pH 8.0).

### Streptavidin purification and on-bead digestion

Lysates were sonicated briefly (2 x 20 sec cycles) and clarified through centrifugation at 20,000 xg for 30 min at 4 °C. Protein concentrations were measured by Bradford and equalized. High-capacity streptavidin sepharose beads (GE healthcare) were washed once with 40 % methanol and twice with ice-cold lysis buffer (1 % sodium deoxycholate in 50 mM ammonium bicarbonate pH 8.0) prior to incubation with protein lysates (50 μL bead slurry per 3 mg total protein) for 2h at 4 °C on a rotator. Beads were then washed twice with cold lysis buffer, and then once with each of the following washing buffers: 1 M KCl, 0.1 M Na_2_CO_3_, 2 M Urea in 50 mM ammonium bicarbonate pH 8.0. The last two washes were performed with cold 50 mM ammonium bicarbonate pH 8.0. Beads were then re-suspended in 2 M Urea in 50 mM ammonium bicarbonate pH 8.0 and protein reduction was performed by incubating with DTT at a final concentration of 10 mM for 45 min at 37 °C and 800 rpm. Samples were subsequently alkylated through incubation with iodoacetamide at a final concentration of 25 mM for 30 min at room temperature and 800 rpm. Samples were pre-digested with 0.5 μg LysC for 4h at 37 °C and 800 rpm and then digested with 1 μg trypsin overnight at 37 °C and 800 rpm. Peptides were acidified to pH 2 by adding formic acid to 1 % final concentration and centrifuged for 15 min at 16,000 rpm at 4 °C to remove any precipitates or beads on Sep-Pak μ-C18 Elution Plates. Peptides were then dried on a speed vac for 2h and re-suspended in 0.1% formic acid prior to LC-MS.

### Mass spectrometry

LC-MS/MS was performed using an Ultimate 3000 RSLCnano (Thermo Fisher Scientific) LC system equipped with an Acclaim PepMap RSLC column (15 cm × 75 μm, C18, 2 μm, 100 Å, Thermo Fisher Scientific). Peptides were eluted at a flow rate of 300 nL/min in a linear gradient of 2-35% solvent B over 70 minutes (solvent A: 0.1% formic acid; B: 0.1% formic acid in acetonitrile (ACN)), followed by a 5 minutes washing step (90% solvent B) and a 10 minutes equilibration step (2% solvent B). Mass spectrometry was performed on a Q-Exactive Plus mass spectrometer (Thermo Fisher Scientific) equipped with a nano-electrospray source and using uncoated SilicaTips (12 cm, 360 μm o.d., 20 μm i.d., 10 μm tip i.d.) for ionization, applying a 1500 V liquid junction voltage at 250°C capillary temperature. MS/MS analysis were performed in data dependent acquisition (DDA) mode. The precursor masses were measured at a resolution of 60,000, and the 12 most intense parent ions were fragmented with a resolving power of 17500 (at 200 m/z). Automatic gain control (AGC), maximum fill time and dynamic exclusion were set to 1e^6^, 60 ms and 30 ms respectively.

### Analysis of time-resolved proteomes

Protein quantification was performed with MaxQuant (version 1.6.17.0) using the following parameters: carbamidomethyl cysteine as fixed modification, methionine oxidation and N-acetylation as variable modifications, digestion mode trypsin specific with maximum 2 missed cleavages, and an initial mass tolerance of 4.5 ppm for precursor ions and 0.5 Da for fragment ions. All experiments were performed in quadruplicate except for the “no peroxide” sample which was performed in triplicate. The abundance of assembled proteins was determined using label-free quantification with the match between runs option was selected. Protein identification was performed searching against the Uniprot reference proteome (Uniprot ID 9615, downloaded on 06 January 2020). Proteins tagged as “Potential contaminants”, “Reverse”, and “Only identified by site” were removed from the analysis. Proteins were matched their respective human gene names as described in Tan et. al., ^43^. A moderated t-test was performed using the Proteomics Toolset for Integrative Data Analysis (ProTIGY, Broad Institute, Proteomics Platform, https://github.com/broadinstitute/protigy). Data was filtered to include only proteins with a minimum of two replicate identifications at each timepoint across the time course dataset. Proteins which were found to be significantly enriched with a p-value less than 0.05 and a log2 fold change greater than 1.0 as compared to both controls were considered for further analysis. 20 proteins (CGN, IRF3, ISG15, KRAS, LGALS9, LMO7, MYO1B, PARD3, PKN2, PPP1R9A, PSME3, PTPN13, RAB35, RPL19, SCRN3, SLC25A1, SLC25A6, TJP1, USP6NL, WASL) which were not found to be significantly enriched, but were identified in our previously published Pals1 proximity proteome ^43^ and satisfied all other criteria were also considered for further analysis. The final proximity proteome encompassed 278 proteins (see Supplementary File S1). Further data analysis and visualization were performed in R (version 4.1.2). Data clustering and heatmap generation were carried out using the ComplexHeatmap package in R. Enrichment analysis was carried out using the Enrichr analysis tool (https://maayanlab.cloud/Enrichr/).

### Generation of interactome

Interaction entries were retrieved from the BioGRID (version 4.3.195) and STRING databases. For BioGRID entries, only physical protein interactions were considered. For STRING entries, only experimental and database interactions with a combined confidence score > 0.4 were included.

### Quantifications and statistical analyses

Immunofluorescence images were analysed in Zeiss Zen software or ImageJ/Fiji ^86,87^. Line scans were performed using the Multichannel plot profile plugin. Pearson’s correlation coefficients were obtained using the Coloc2 plugin by marking the junctions as regions of interest. For other measurements, such as length and area, the “measure functions” in ImageJ were used. For the measurement of apical F-actin intensity, cell boundaries were segmented using Cellpose ^92^ and apical F-actin intensity measured using maximum intensity projections of the apical portion of the z-stack. Quantification of junctional Pals1 levels was performed on maximum intensity projections of z-stacks using scikit-image (segmentation), NumPy/SciPy (statistics), and Matplotlib (visualisation). In brief, ZO-1 signals were thresholded, binarized, and skeletonized to a 1-pixel centreline, which was dilated by 3 pixels. This mask was used to measure the mean junctional fluorescence intensities of Pals1 and PatJ per cell. Values are represented as mean ± SEM unless stated otherwise. Statistical analyses were performed using a two-tailed unpaired Student’s t-test in GraphPad Prism unless stated otherwise. All data represent at least three independent repeats: n.s, not significant, *p<0.05, **p<0.01, ***p<0.001, ****p<0.0001.

