## Supplementary Figures for "Bulk delivery of a preassembled apical surface initiates epithelial lumen formation"

**Supplementary information and files**

**Supplementary File S1:** Mass spectrometry data of the 278 proteins identified by time-resolved Pals1-APEX2 proximity labeling, related to Figure 3

**Supplementary File S2:** Materials and Resources

**Supplementary Movies:**

**Movie S1:** FIB-SEM of an intracellular VAC, related to Figure 6A
**Movie S2:** 3D segmentation of microvilli in an intracellular VAC, related to Figure 6A
**Movie S3:** FIB-SEM of a microvilli-containing membrane pit at the ECM-facing surface, related to Figure 6B
**Movie S4:** 3D segmentation of peripheral ER and protrusions on the ECM-facing surface, related to Figure 6C

**Supplementary Figures S1-S15**

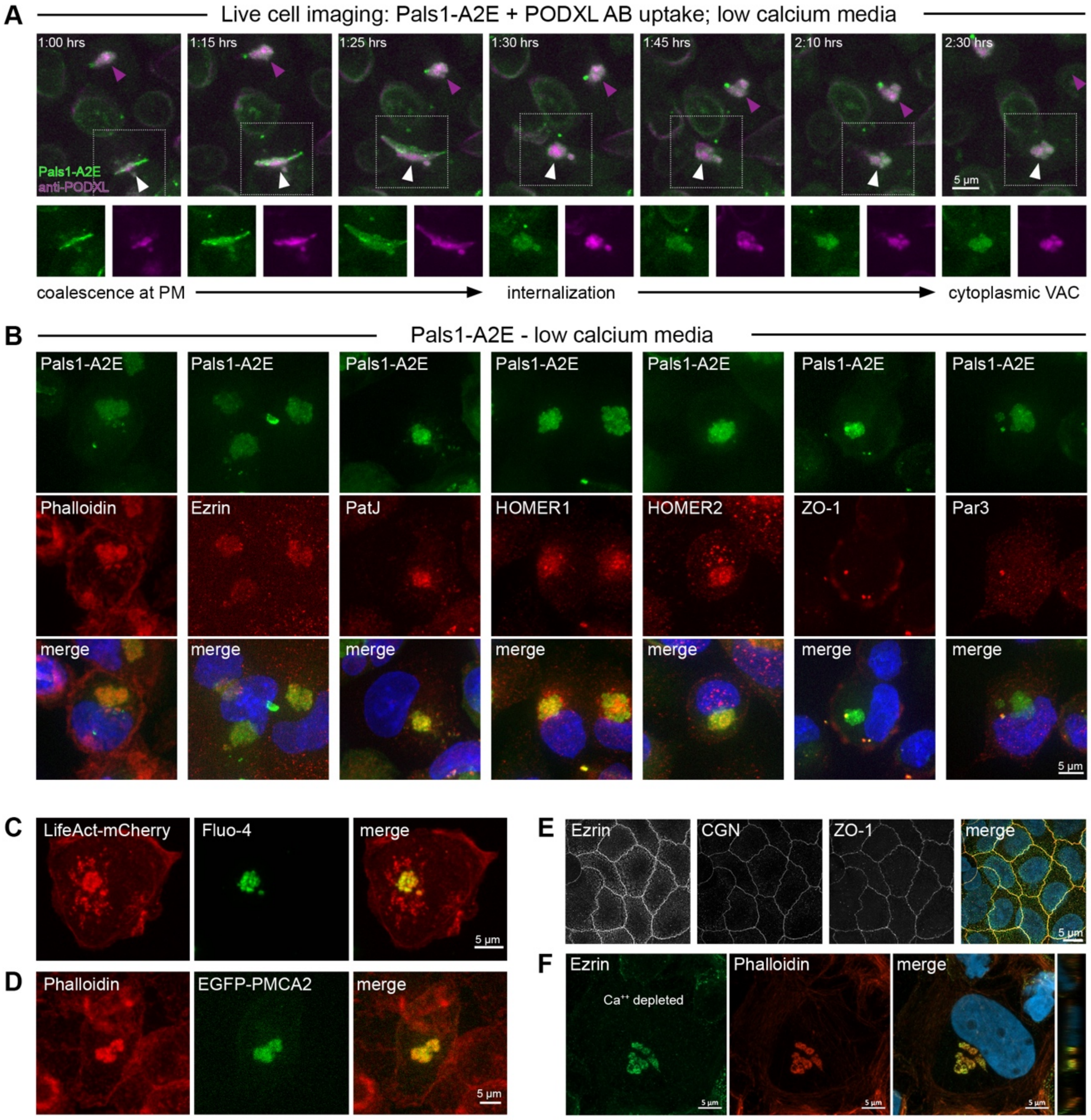


**Figure S1: VAC formation and composition**

(A) Time-lapse imaging of MDCK cells stably expressing Pals1-A2E incubated with anti-PODXL antibody. Pals1 and PODXL antibody are co-internalized into large organelles (VACs) in response to calcium depletion. White arrowheads indicate an internalization event leading to VAC formation, magenta arrowheads point to an existing VAC.

(B) Pals1-A2E MDCK cells were calcium-depleted overnight, fixed and stained with the F-actin dye phalloidin or the indicated antibodies.

(C and D) MDCK cells transfected with LifeAct-mCherry (C) or EGFP-PMCA2-wb (D) were grown in low-calcium medium overnight. The calcium binding dye Fluo-4 was added to live cells transfected with LifeAct-mCherry. EGFP-PMCA2-wb transfected cells were fixed and stained with phalloidin.

(E) Caco-2 monolayers were fixed and stained with antibodies against ezrin, CGN and ZO-1.

(F) Caco-2 monolayers were cultured in calcium-free medium overnight, fixed and stained with anti-ezrin antibodies and phalloidin.

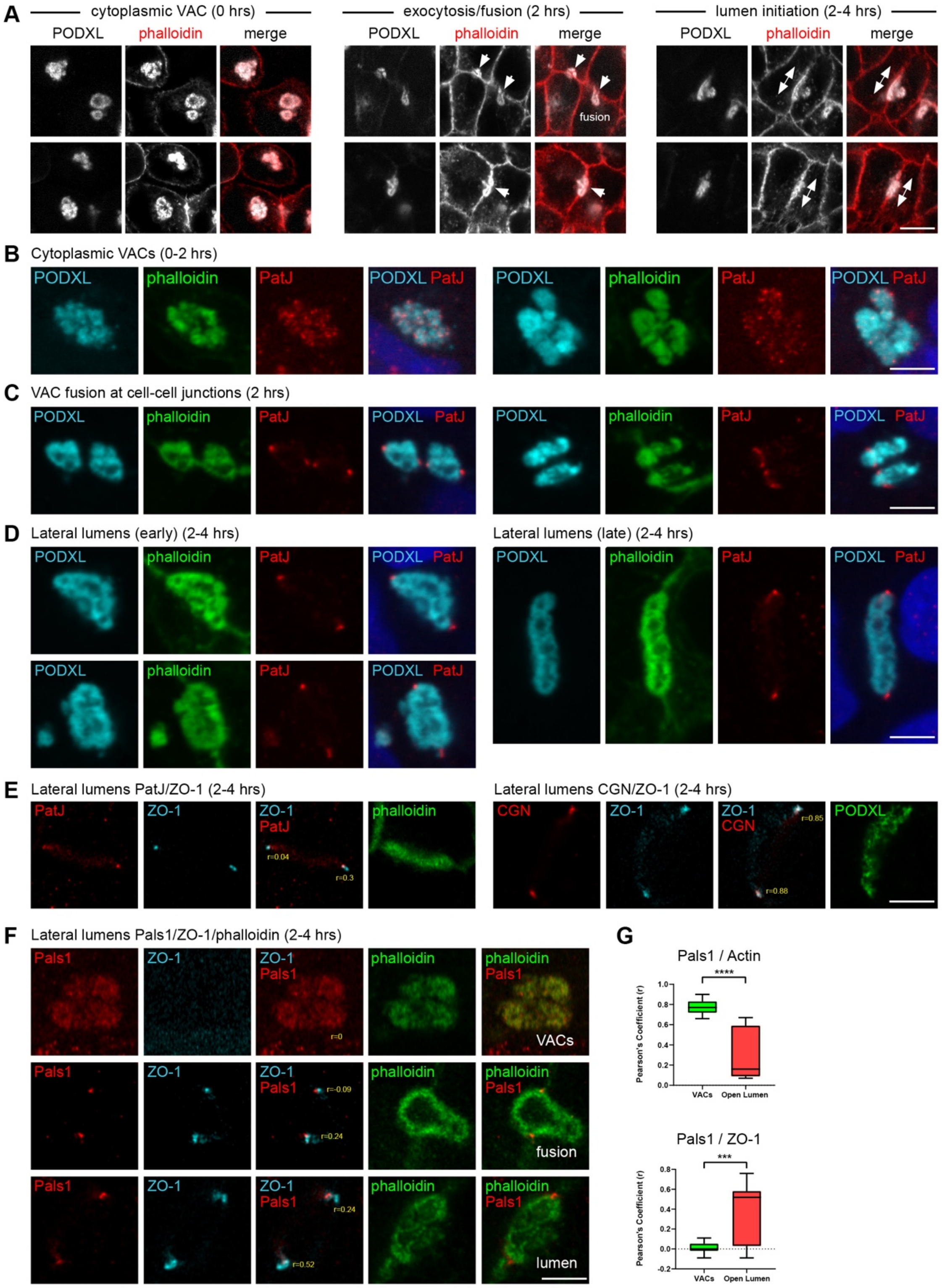


**Figure S2: TJ and VMZ proteins form a junctional belt around lateral lumens**
(A) MDCK cells were fixed 0-4h post calcium addition and stained with anti-PODXL antibodies and phalloidin. Intracellular VACs (left), VAC fusion sites (middle), and lateral lumens (right) are shown. Scale bar 5 µm.
(B-D) MDCK cells were fixed 0-4h post calcium addition and stained with anti-PODXL and anti-PatJ antibodies, and phalloidin. Intracellular VACs (B), VAC fusion sites (C), and lateral lumens (D) are shown. Scale bar 5 µm.

(E) MDCK cells were fixed 2-4h post calcium addition and stained with anti-ZO-1 and anti-PatJ antibodies, or anti-ZO-1 and anti-CGN antibodies. Lateral lumens are shown. Pearson’s correlation coefficients (r) are shown. Scale bar 5 μm.
(F) MDCK cells were fixed 0-4h post calcium addition and stained with anti-ZO-1 and anti-Pals1 antibodies. Intracellular VACs, VAC fusion sites, and lateral lumens are shown. Pearson’s correlation coefficients (r) are shown. Scale bar 5 μm.
(G) Pearson’s correlation coefficients of Pals1 and actin (phalloidin) and Pals1 and ZO-1 markers in VACs and at lateral lumens.

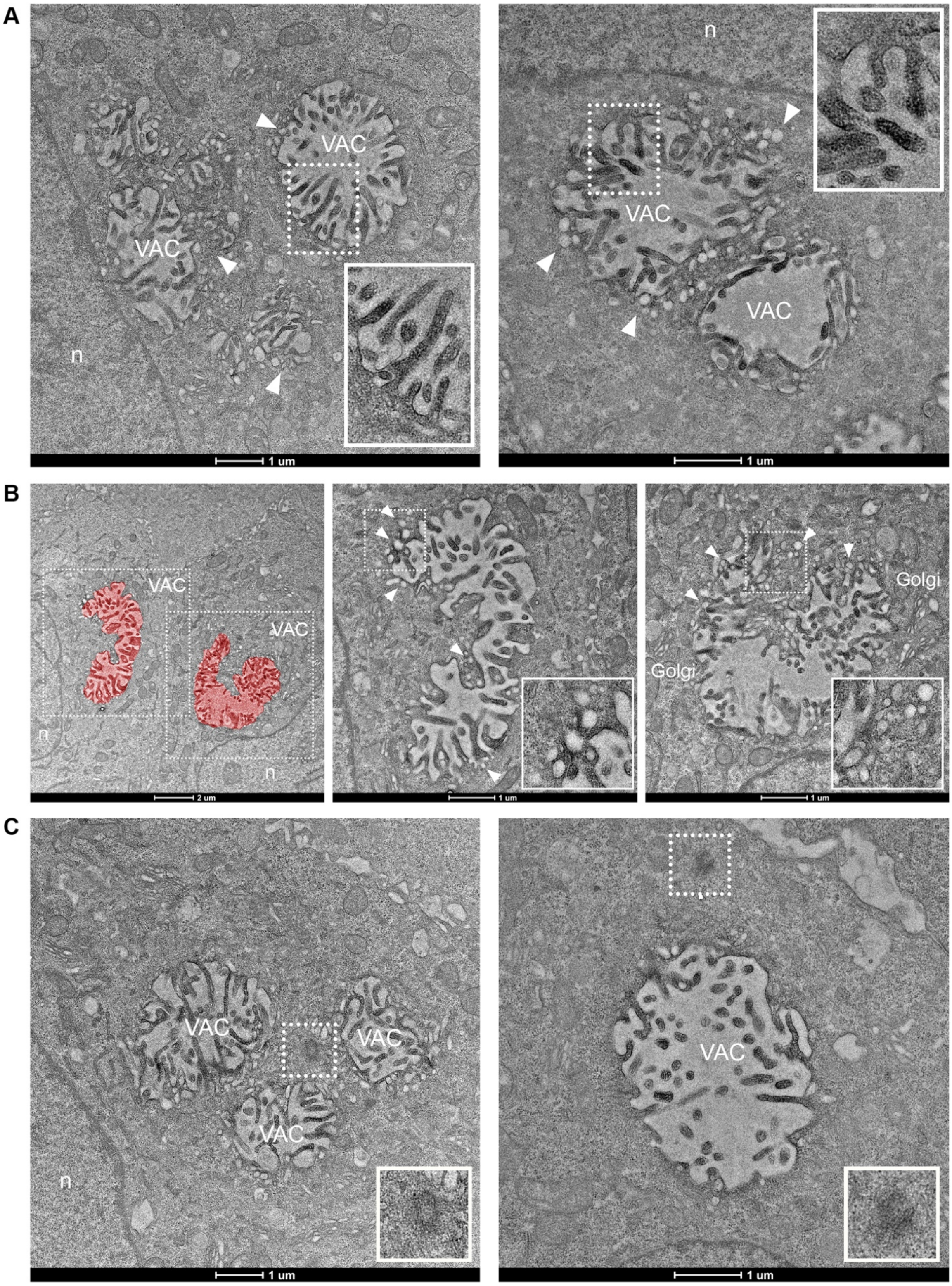


**Figure S3: Ultrastructure of VACs**
(A-C) Representative TEM micrographs of calcium-depleted Pals1-A2E MDCK cells. Note the strong EM contrast in microvilli facing the VAC lumen (A), and the presence of vesicles surrounding the VACs (B; white arrowheads). Centrioles are also frequently observed close to VACs (C, insets). n = nucleus.

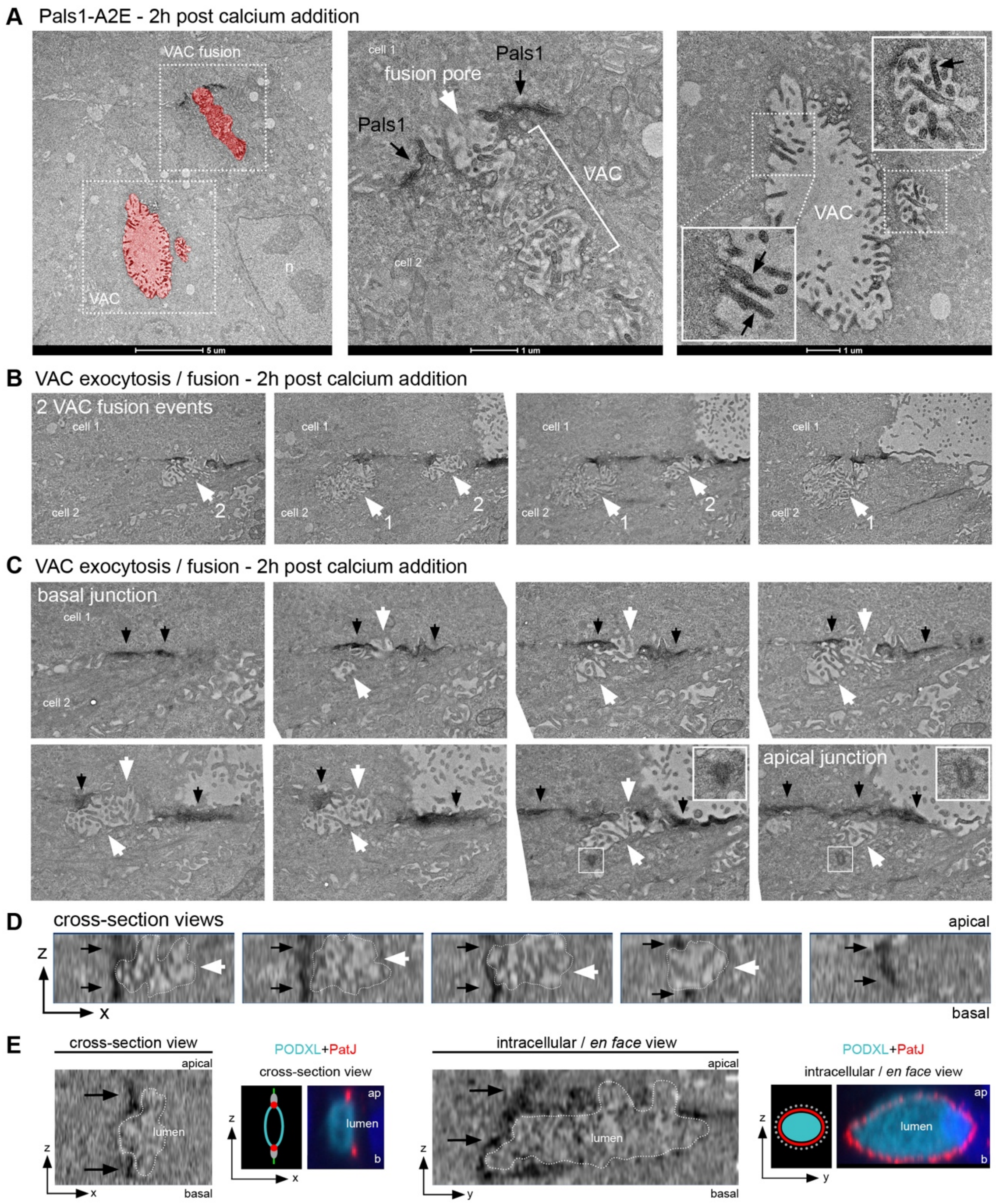


**Figure S4: Ultrastructural analysis of VAC exocytosis**
(A) Representative TEM micrographs of Pals1-A2E MDCK cells 2h after calcium addition. An intracellular VAC and a VAC fusion site at the lateral membrane (corresponding to data shown in Figure 2B) are shown. Note the increase in EM contrast at cell junctions surrounding the VAC fusion site and in apical microvilli (black arrowheads). n=nucleus.
(B) Serial section TEM analysis of two adjacent VAC fusion sites at the lateral membrane. Select serial sections are shown. Fusion site 1 relates to data shown in Figure 2C. Fusion site 2 is shown in detail in (C).
(C) Serial section TEM of VAC fusion shown in (B). Note the increase in junctional EM contrast produced by Pals1-A2E (black arrowhead) around the VAC fusion site (white arrowheads). Centrioles are localised in close proximity to the VAC (insets).

(D) Cross-section views of a VAC fusion site (related to Figure 2C). The lumen is outlined by white dots. Note that Pals1-A2E (black arrows) forms a junctional ring around the lumen.
(E) Cross-section and *en face* views of a lateral lumen (related to Figure 2D). The lumen is outlined by white dots. Note that Pals1-A2E (black arrows) forms a junctional ring around the lumen. Corresponding projections of confocal z-stacks of a lateral lumen are shown. Cells were stained with antibodies against PODXL and PatJ.

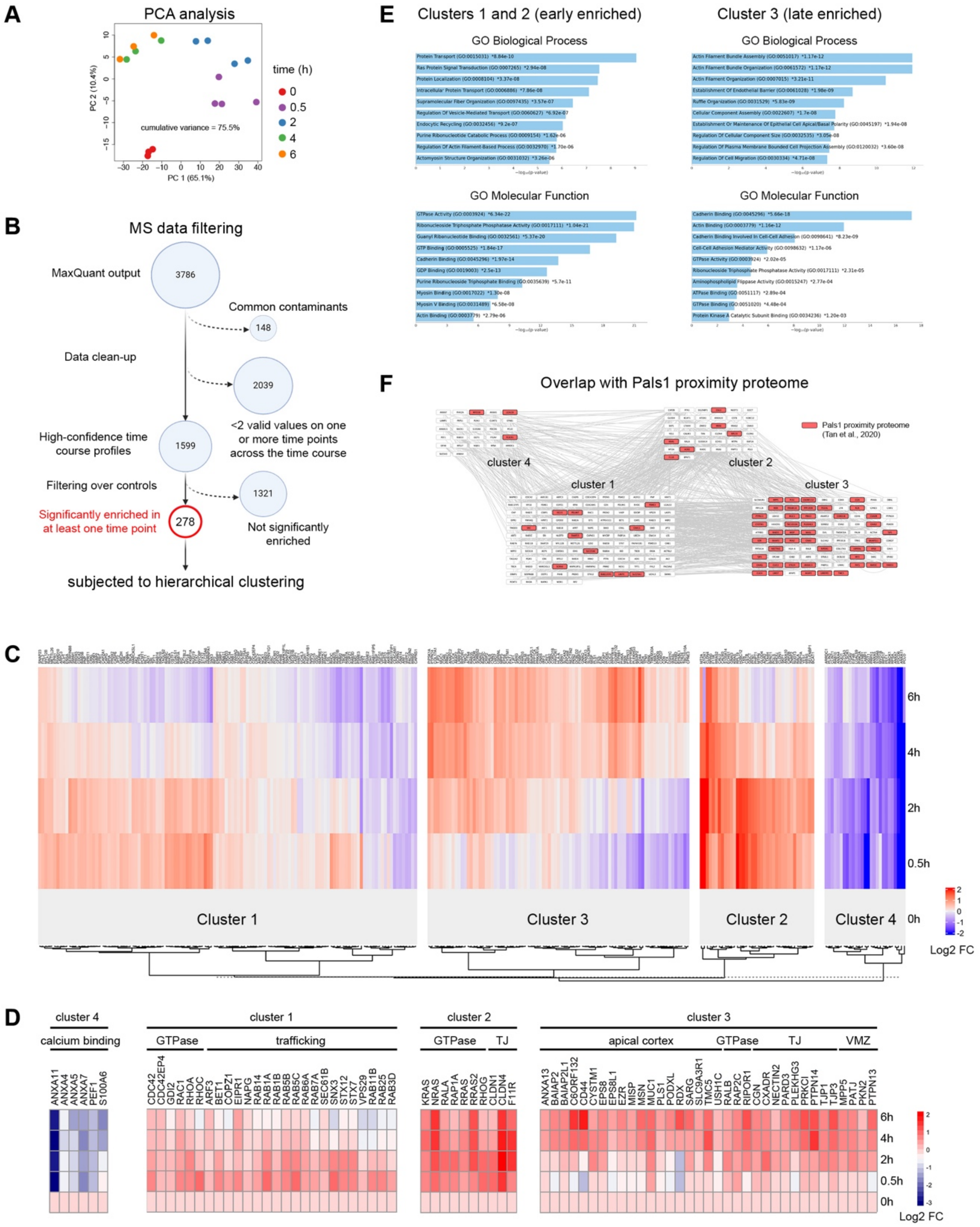


**Figure S5: A Time-resolved proximity proteome of Pals1 during cell junction assembly**

(A) Principal Component Analysis (PCA) of the proteomics data.

(B) Flow chart summarizing the LC-MS/MS analysis and proteomics data filtering pipeline. Label-Free Quantification (LFQ) was used to define proteins significantly enriched as compared to the controls. 278 proteins were significantly enriched in at least one time point and subjected to hierarchical clustering.

(C) Dendrogram and heatmap generated from hierarchical clustering of the 278 identified proteins. Dotted line indicates where the dendrogram was cut to generate four clusters. Heat map shows the log2-fold changes relative to the 0h timepoint.

(D) Heatmap showing the log2 fold changes of select protein categories from each cluster.
(E) Gene Ontology (GO) annotations of clusters 1+2 and cluster 3.
(F) Interaction network of the 278 identified proteins arranged by cluster. Interaction network is based on STRING and BioGRID entries. The 65 proteins highlighted in red were previously identified in a static Pals1 proteome generated from fully mature epithelial monolayers [1]. Note that more than two thirds of these (45 out of 65) are enriched in cluster 3.

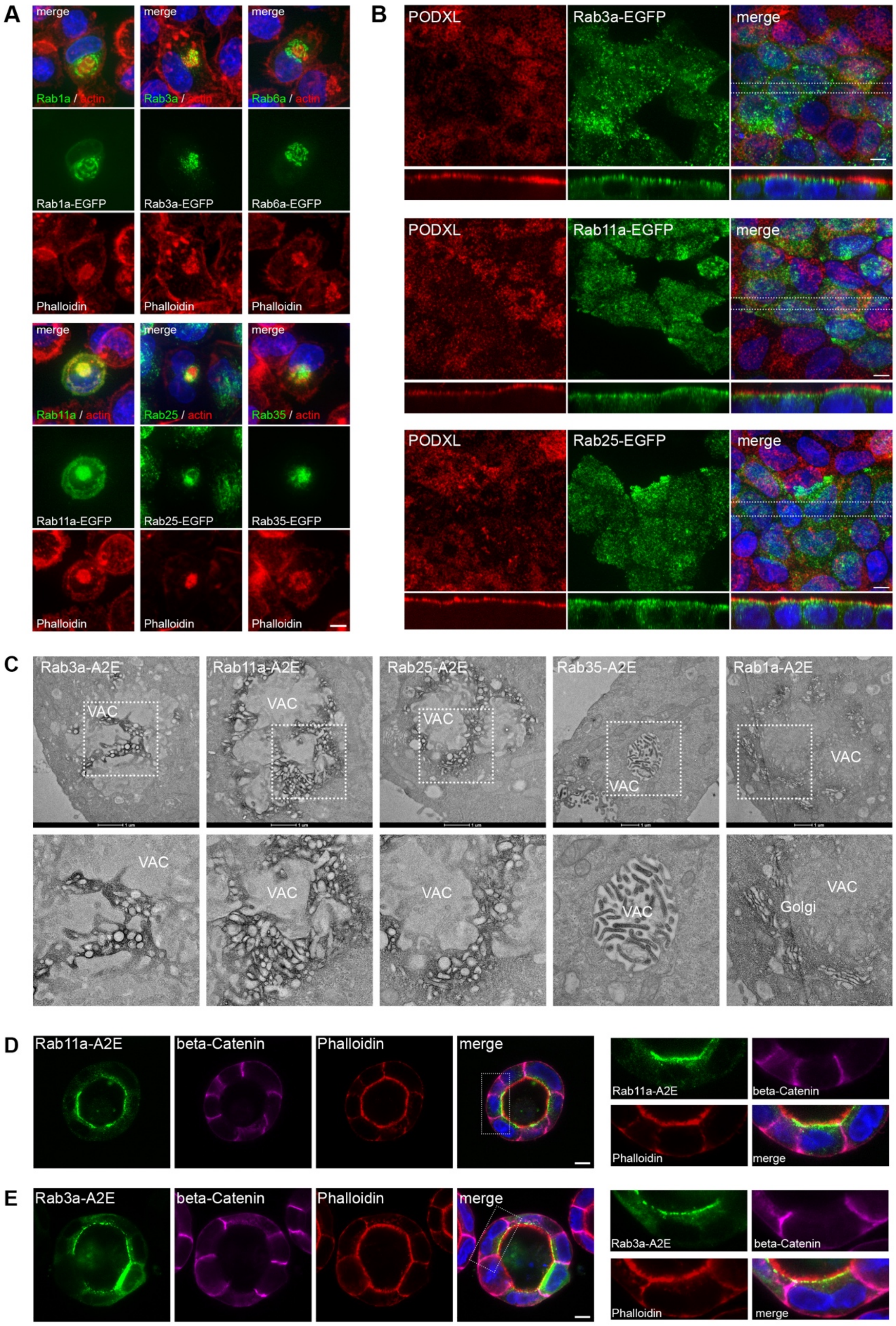


**Figure S6: Apical Rab GTPases are associated with VACs**
(A) MDCK cells stably transfected with EGFP-tagged Rab proteins were calcium-depleted overnight, fixed, and co-stained with phalloidin. All Rab proteins analysed are associated with VACs. Scale bar 5 µm.

(B) MDCK cells stably transfected with EGFP-tagged Rab proteins were cultured on Transwell filters for 10 days, fixed and co-stained with anti-PODXL antibodies. Confocal x/y projections and x/z projections (of subvolumes indicated in the merged image) are shown. Note that all Rabs analysed localize apically. Scale bar 5 µm.
(C) MDCK cells stably transfected with Rab-APEX2-EGFP proteins were calcium-depleted overnight and processed for APEX2-TEM. Representative micrographs are shown. Rab3, Rab11, and Rab25 are associated with small vesicles close to the VAC, whilst Rab35 localises specifically to VAC microvilli. Rab1 localises to the Golgi and is not associated with VACs.
(D and E) MDCK cells stably transfected with Rab11-APEX2-EGFP (D) or Rab3-APEX2-EGFP (E) were grown in Matrigel for 4 days, fixed and stained with phalloidin and antibodies against beta-catenin. Representative confocal micrographs of end-stage 3D cysts are shown. Scale bar 10 µm.

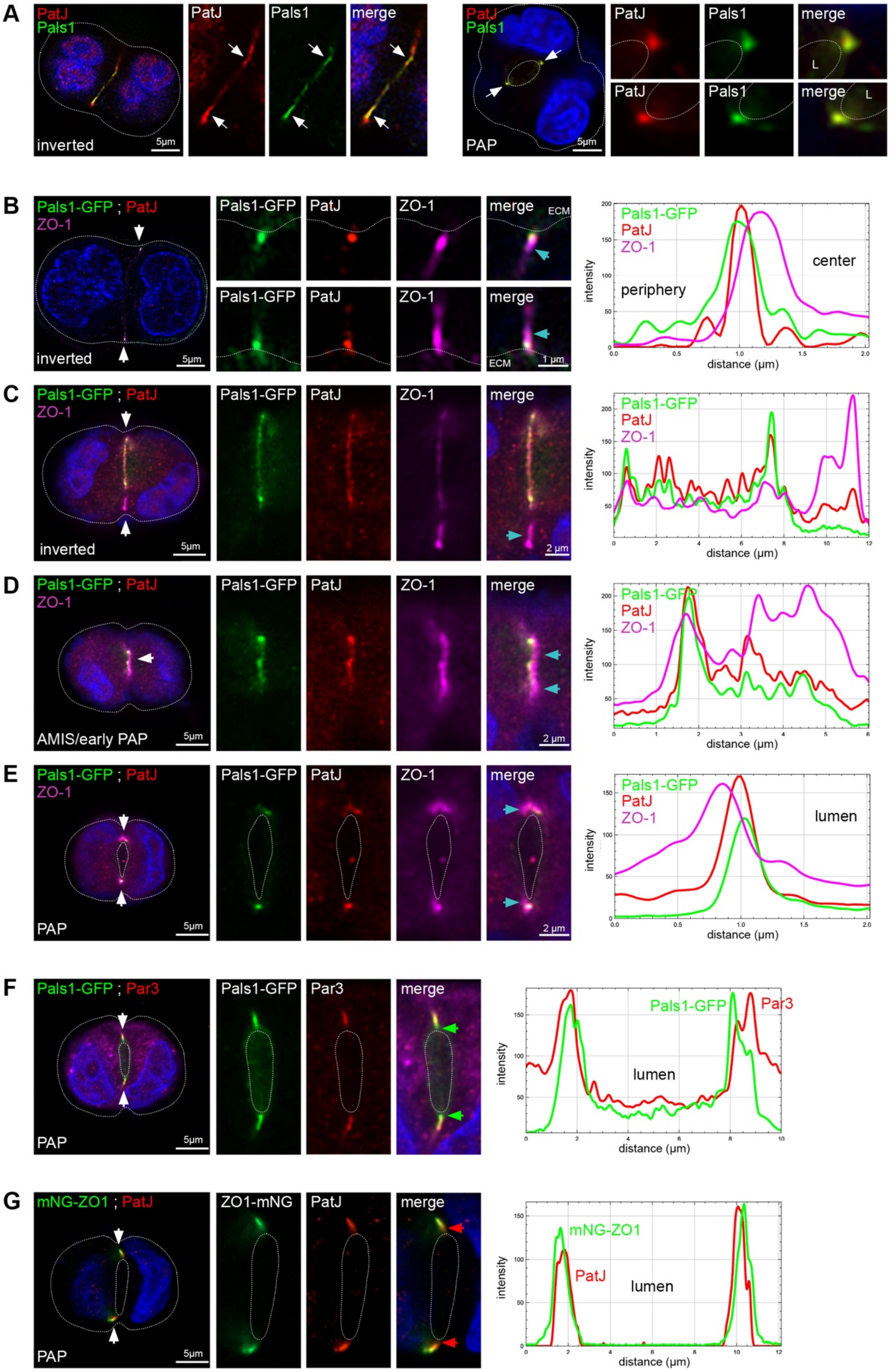


**Figure S7: Spatio-temporal dynamics of TJ and VMZ proteins during lumen initiation**

(A) Inverted and PAP stages of wild-type MDCK cells stained with antibodies against Pals1 and PatJ.
(B-E) Inverted (B and C), AMIS/early PAP (D), and PAP (E) stages of MDCK cells stably transfected with Pals1-GFP stained with antibodies against PatJ and ZO-1. Cyan arrowheads demarcate ZO-1. Line scans illustrate colocalisation of Pals1 and PatJ and the spatial offset between Pals1/PatJ and ZO-1.
(F) PAP stage of MDCK cells stably transfected with Pals1-GFP stained with anti-Par3 antibodies. Green arroweads demarcate Pals1. Line scans illustrate the spatial offset between Pals1 and Par3.
(G) PAP stage of genome-engineered MDCK cells expressing endogenously tagged mNeonGreen-ZO-1 stained with anti-PatJ antibodies. Red arroweads demarcate PatJ. Line scans illustrate the spatial offset between PatJ and ZO-1.

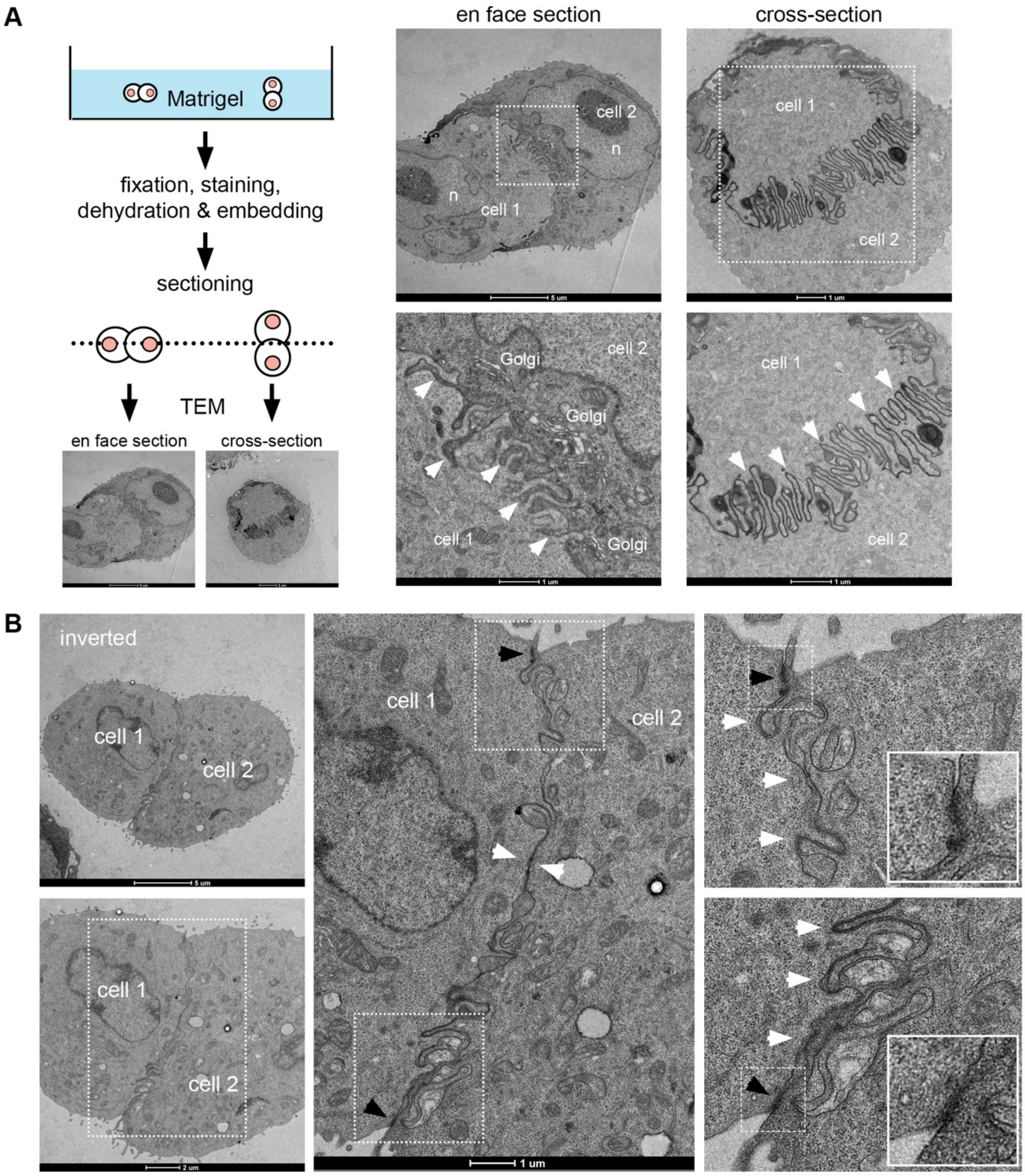


**Figure S8: Ultrastructure of early-stage cell doublets embedded in Matrigel**
(A) MDCK cells grown for 18h in Matrigel were processed for TEM. Representative *en face* and cross-sections through cell doublets are shown. Note the abundance of interdigitated cell junctions at the cell-cell contact site (white arrowheads).
(B) Representative *en face* micrographs showing interdigitated cell junctions (white arrowheads) and electron-dense cell junctions at the ECM-facing end of the cell-cell contact interface (black arrowheads).

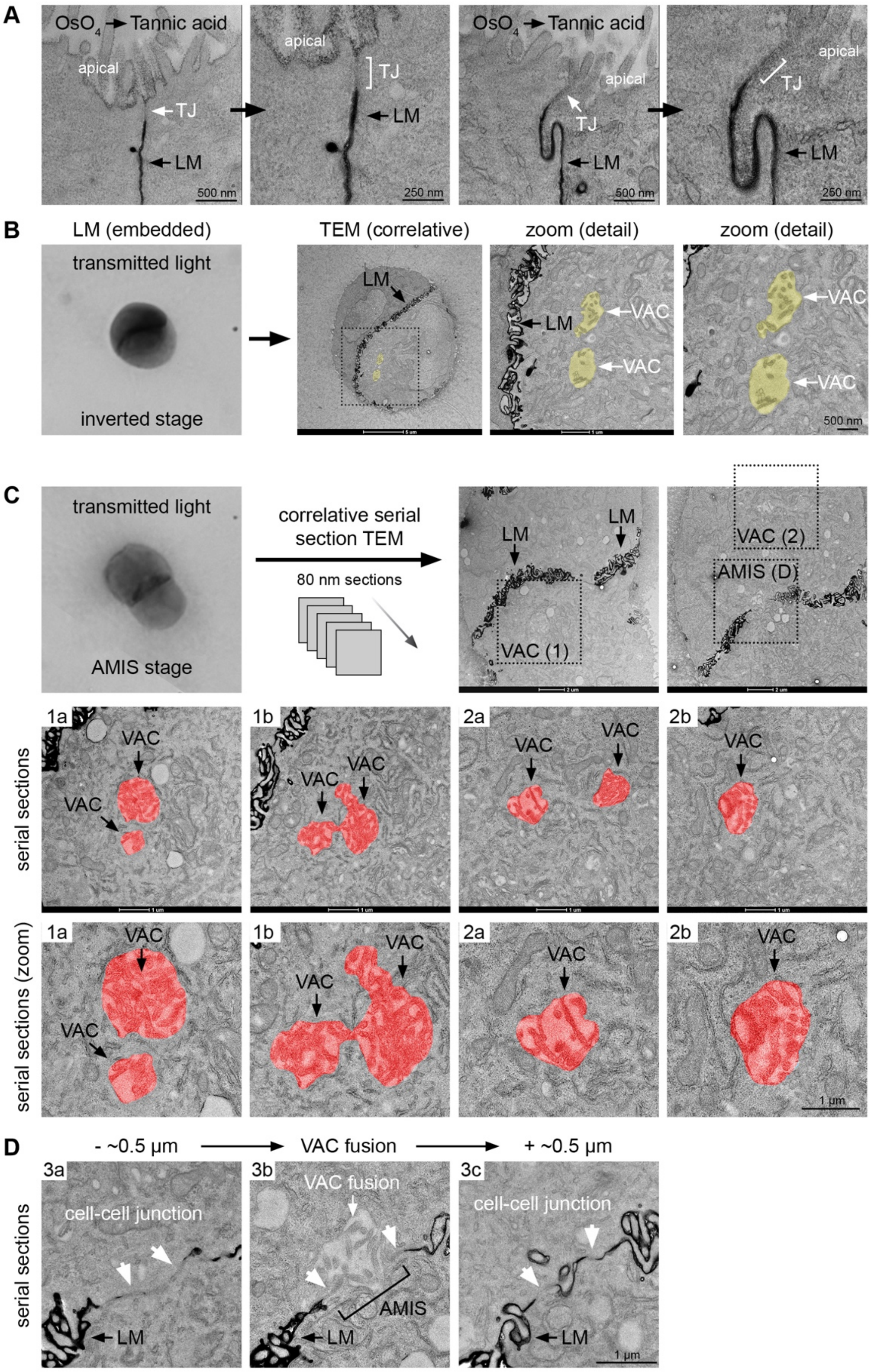


**Figure S9: Serial section TEM of VACs and VAC fusion at the AMIS**
(A) Two representative TEM micrographs of MDCK monolayers processed with the osmium/tannic acid staining protocol. Note the strong EM contrast at the lateral membrane (LM). Tight junctions (TJ) appear electron-lucent, and the apical membrane is unstained.

(B) MDCK cells embedded in Matrigel for 24h were processed for correlative light and serial section TEM using the osmium/tannic acid staining protocol as in (A). Transmitted light images and representative correlative TEM micrographs of an inverted stage are shown. Intracellular VACs are highlighted.
(C) MDCK cells embedded in Matrigel for 24h were processed for correlative light and serial section TEM using the osmium/tannic acid staining protocol as in (A). Transmitted light images and representative serial section TEM micrographs showing VACs at the AMIS stage are shown. Related to Figure 5F.
(D) Serial section TEM of a VAC fusion event at the AMIS (corresponding to the area indicated as (D) in panel (C)). The fusion site (middle panel) and micrographs 0.5 µm below and above the fusion site are shown. Related to Figure 5F.

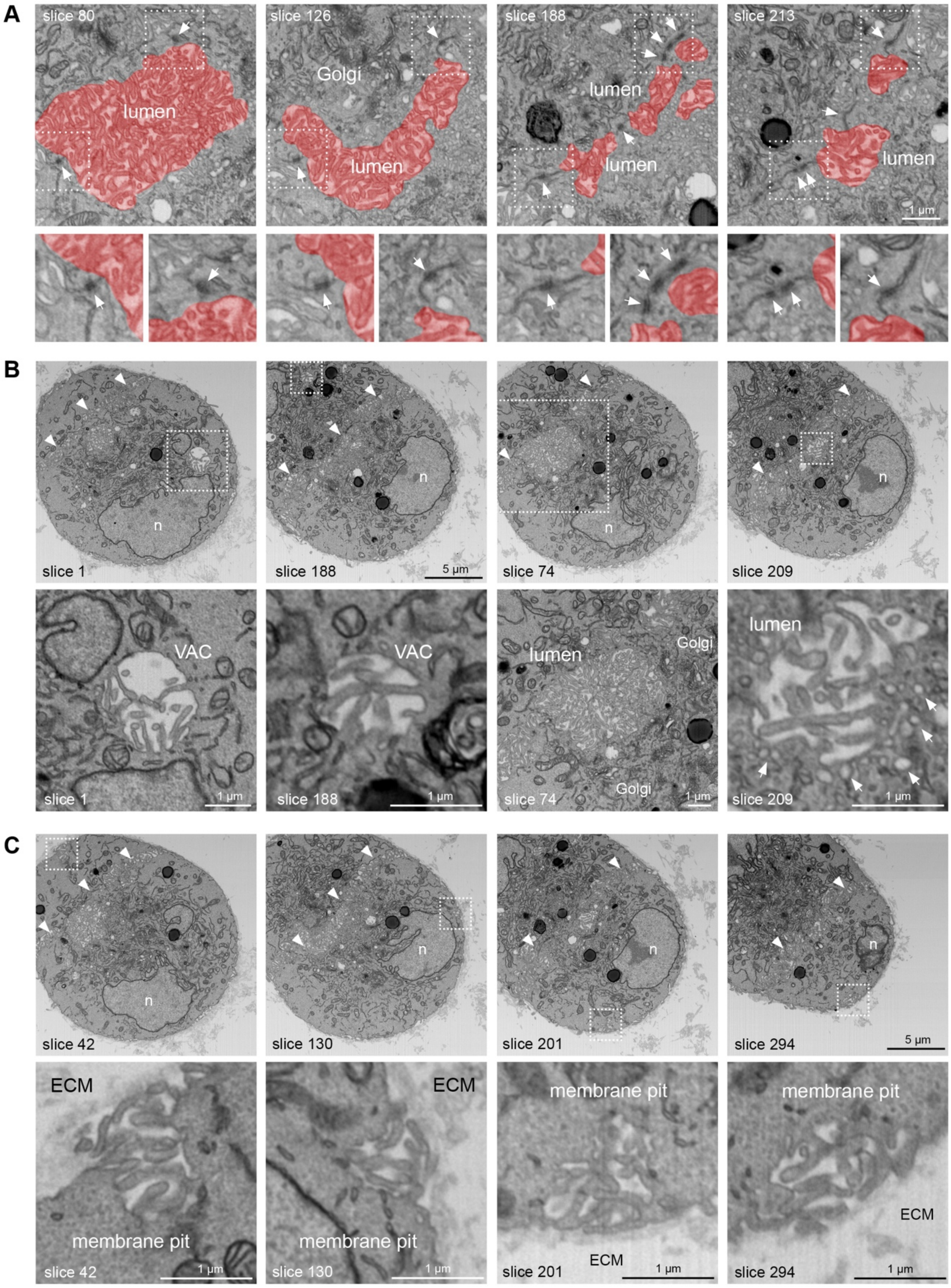


**Figure S10: Ultrastructure of the PAP stage analysed by FIB-SEM**
(A) Representative slices of the FIB-SEM volume showing cell-cell junctions (white arrows) surrounding individual apical lumens.
(B) Representative slices of the FIB-SEM volume showing two VACs and two apical lumens. Note the presence of vesicles (white arrowheads) and Golgi cisternae close to the apical luminal surface.

(C) Representative slices of the FIB-SEM volume showing four microvilli-rich membrane pits at the ECM-facing plasma membrane.

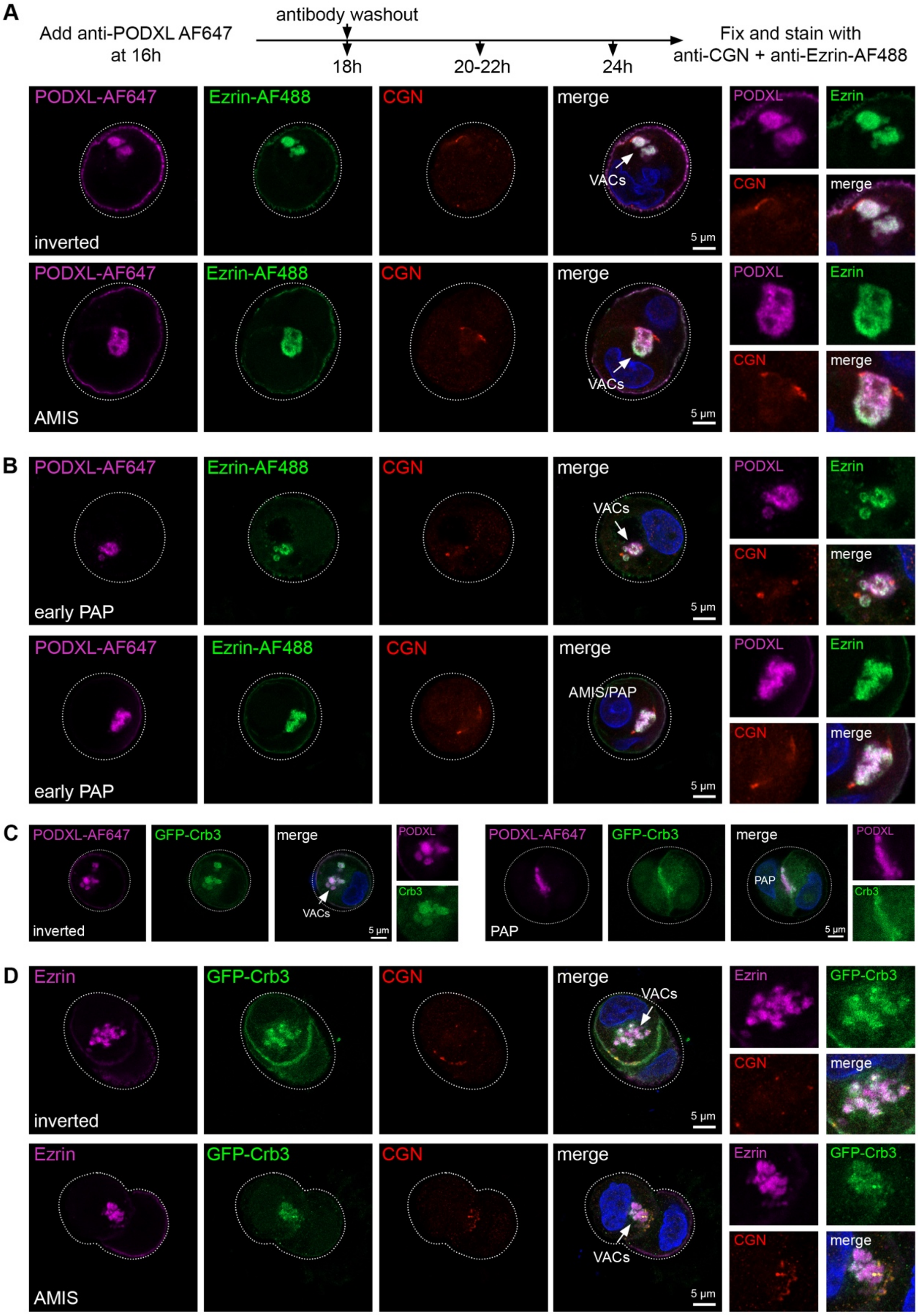


**Figure S11: VACs deliver apical proteins to the AMIS via transcytosis**
(A and B) Transcytosis of PODXL was analysed using an antibody uptake assay. AF647-labelled PODXL antibody was added at 16h and washed out at 18h. Cells were fixed between 18h and 24h and stained with anti-CGN antibodies and a pre-labeled Ezrin-AF488 antibody. Representative inverted, AMIS, and early PAP stages are shown. Related to Figure 6F.

(C) PODXL antibody uptake assay as in (A) in MDCK cells stably transfected with GFP-Crb3. Cells were fixed between 18h and 24h. Representative inverted and PAP stages are shown. Related to Figure 6G.
(D) MDCK cells stably transfected with GFP-Crb3 were fixed between 18h and 24h and stained with anti-Ezrin and anti-CGN antibodies. Representative inverted and AMIS stages are shown. Related to Figure 6H.

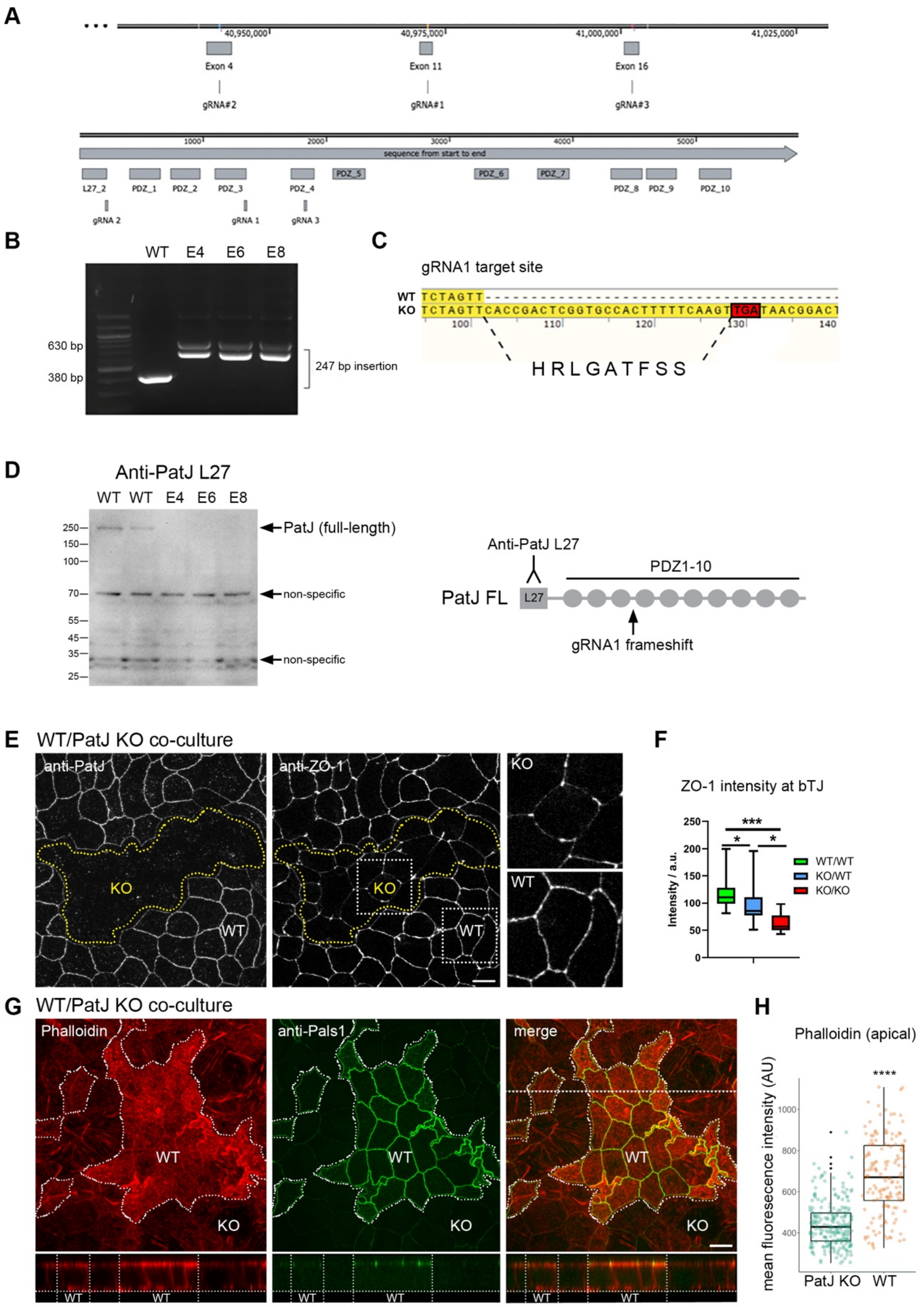


**Figure S12: Generation and characterisation of PatJ KO MDCK cells**
(A) Graphical illustration of the canine *PatJ* gene locus. The target sites of the three sgRNAs used are shown.
(B) PCR analysis of the gene locus surrounding the sgRNA1 target site in MDCK WT cells and PatJ KO clones E4, E6 and E8. DNA sequencing of the PCR products revealed a 247 bp insertion in all three KO clones.

(C) DNA sequence of the sgRNA1 target site in PatJ KO clone E4. The 247 bp insertion results in a Stop codon 9 amino acids downstream of the sgRNA target site.
(D) Western blot analysis of PatJ KO clones E4, E6, and E8 using an antibody against the L27 domain of PatJ. Note the complete loss of PatJ expression.
(E) MDCK WT cells were cocultured with PatJ KO cells on Transwell filters for 12 days, fixed and stained with anti-PatJ and anti-ZO-1 antibodies.
(F) Quantification of bicellular ZO-1 levels based on data shown in (E). ***$p<0.001$, *$p<0.05$; Student's t-test (n=11 junctions/condition). Scale bar 10 µm.
(G) MDCK WT cells and PatJ KO cells were co-cultured on Transwell filters for 12 days. Cells were fixed and stained with anti-PatJ antibodies and phalloidin. x/y and x/z slices of confocal micrographs are shown. Scale bar 10 µm.
(H) Quantification of data shown in (G) using maximum intensity projections generated from the apical portion of the z-stack. **** $p<0.0001$; Student's t-test (PatJ KO, n=226 cells; WT, n=152 cells).

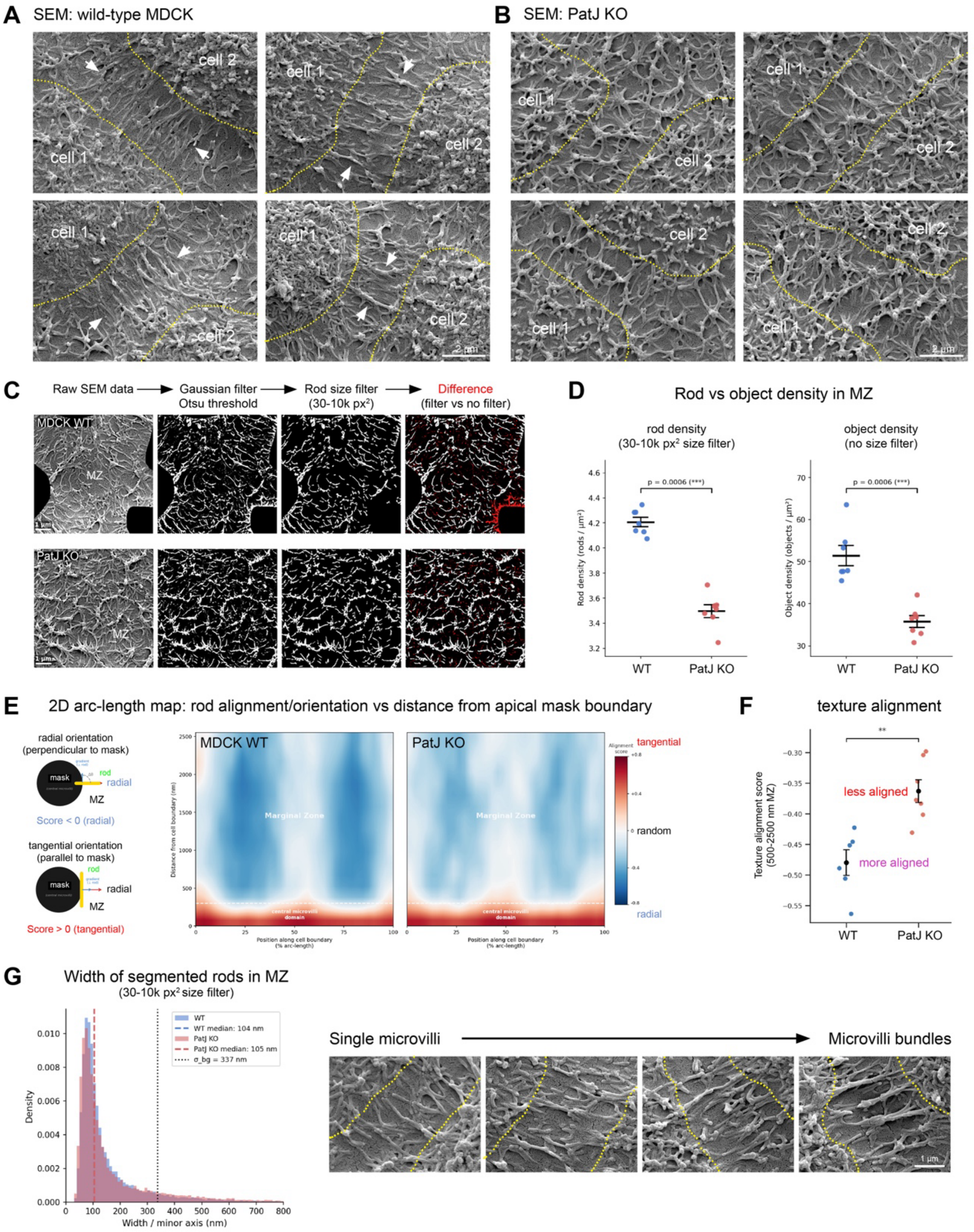


**Figure S13: Automated segmentation and quantification of marginal zone rods**
(A and B) Representative SEM micrographs of the marginal zone (MZ) of MDCK WT (A) and PatJ KO (B) cells cultured on Transwell filters for 16 days. The yellow dotted lines indicate the boundary between the central microvilli-rich domain and the MZ. Parallel arrays of trans-junctional rods are highlighted by white arrowheads. In PatJ KO cells, the MZ appears disorganized.
(C) Workflow depicting the segmentation and quantification of apical rods in the marginal zone (MZ). The central microvilli-rich domain was masked and excluded from the analysis. Rods in the MZ were segmented by applying an area/size filter with a 30-10,000 px$^2$ range.

(D) Direct comparison of rod and object density in the MZ. Left: Rod density determined from thresholded micrographs using a size filter of 30-10,000 $px^2$. Right: Object density determined from thresholded micrographs without applying a size filter. Both quantification approaches reveal a reduced density of microvilli-like structures in the MZ of PatJ KO cells (related to Figure 7F). Statistical analysis was performed using a two-sided Mann-Whitney U test. *** $p<0.001$ (n=6-7 micrographs/condition; 30-40 cells/condition).
(E) 2D arc-length map illustrating the orientation of objects (rods) in the MZ as a function of their distance from the central microvilli-rich domain (black mask). Note that rods within 400-500 nm from the mask boundary (white dashed horizontal lines) are oriented tangentially (red), whilst those within the marginal zone (~500-2500 nm) are oriented radially (blue).
(F) The texture alignment (orientational coherence) of rods in the 500-2500 nm MZ band was quantified (related to Figure 7F). A negative score indicates high orientational order, while a score nearer to zero indicates a more random orientation. Error bars represent mean ± SEM. Statistical analysis was performed using a two-sided Mann-Whitney U test. **$p<0.01$ (n=6-7 micrographs/condition; 30-40 cells/condition).
(G) Width of rods in the MZ of WT and PatJ KO cells measured with a 30-10,000 $px^2$ size filter. Note that rod widths (~105 nm) are identical in WT and PatJ KO cells. Representative high resolution SEM micrographs of microvilli rods in the marginal zone of WT cells are shown.

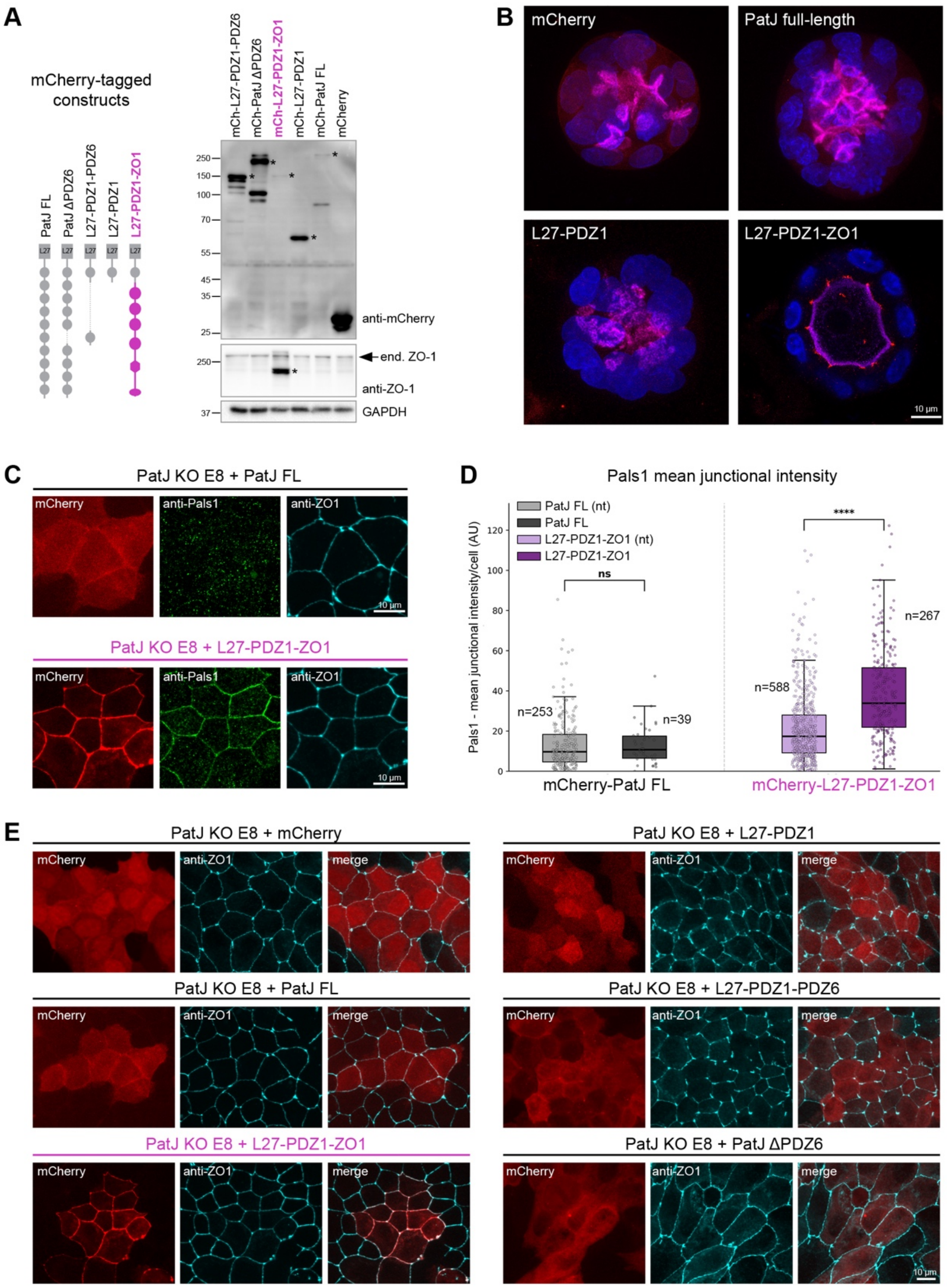


**Figure S14: Rescue experiments in PatJ KO cells**
(A) Western blot analysis of PatJ KO MDCK cells stably transfected with the indicated PatJ constructs and a chimeric PatJ L27-PDZ1-ZO1 construct. All constructs were expressed as N-terminal mCherry fusion proteins. The expected molecular weight of the full-length proteins is indicated with an asterisk (*).
(B) Representative end-stage cysts (4 days in Matrigel) of PatJ KO rescue cell lines. mCherry fluorescence is shown in red, PODXL staining is shown in magenta. Note that only the chimeric PatJ L27-PDZ1-ZO1 construct localises to TJs and rescues the PatJ multi-lumen phenotype.

(C) Confocal micrographs of PatJ KO cells stably transfected with either full-length PatJ or the L27-PDZ1-ZO1 chimera. Cells were stained with anti-Pals1 and anti-ZO-1 antibodies. Note that the L27-PDZ1-ZO1 chimera but not full-length PatJ restores Pals1 recruitment to apical cell junctions.
(D) Junctional Pals1 intensity was quantified in PatJ KO cells stably transfected with either full-length PatJ or the L27-PDZ1-ZO1 chimera. For each cell line, junctional Pals1 levels were measured in mCherry positive and mCherry negative cells (non-transfected, nt). Note that expression of the L27-PDZ1-ZO1 chimera but not full-length PatJ significantly increases junctional Pals1 levels. ****$p<0.0001$, Mann-Whitney U test (n=292 and 855 cells/condition).
(E) Confocal micrographs of PatJ KO cells stably transfected with the indicated mCherry-tagged PatJ constructs. Cells were fixed and stained with ZO-1 antibodies. Note that none of the constructs except the L27-PDZ1-ZO1 chimera are efficiently recruited to apical cell junctions.

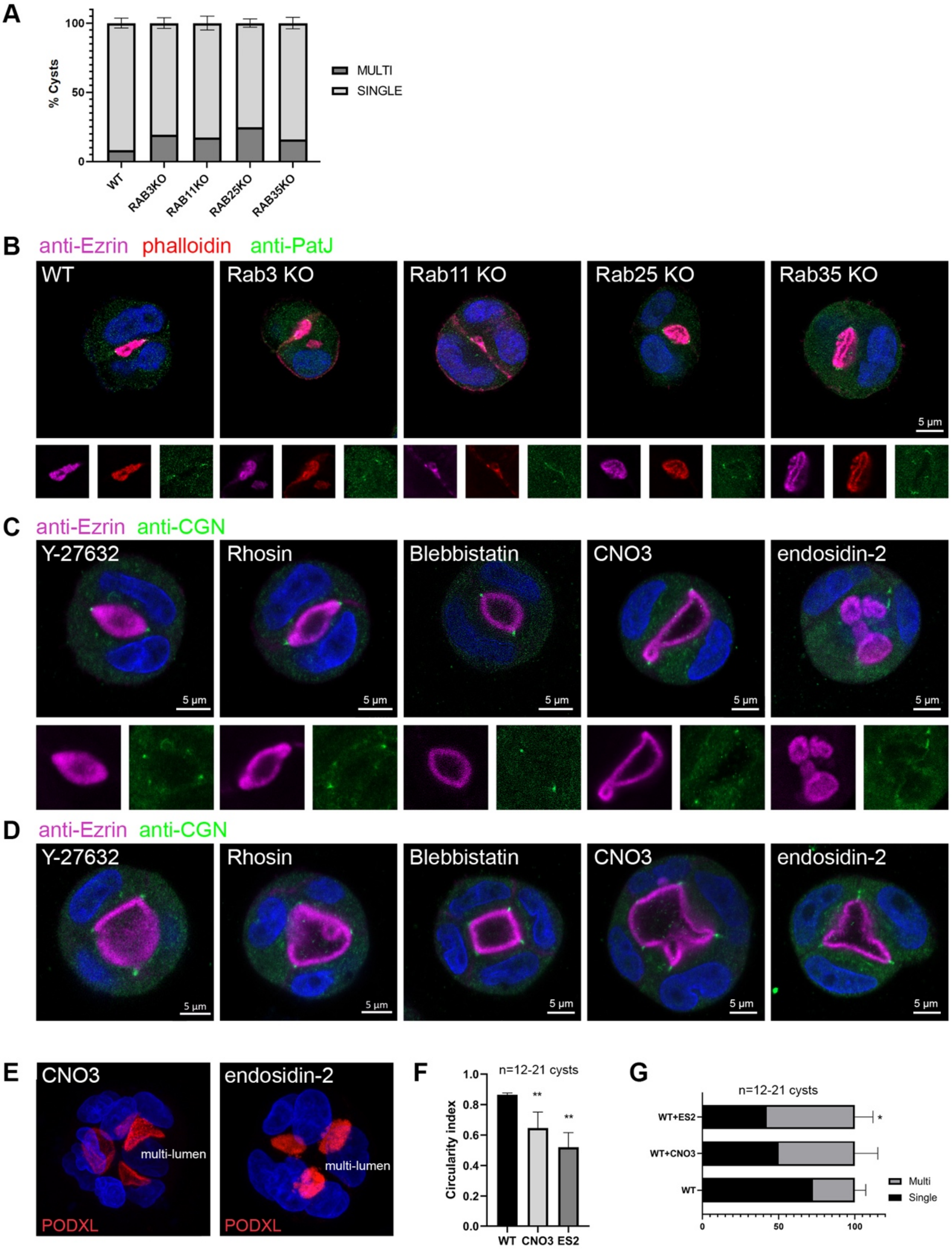


**Figure S15: Function of apical Rab proteins and RhoA/actomyosin activity in lumen formation**

(A) Quantification of cyst morphology in WT MDCK cells and Rab3 (Rab3a/b/c/d quadruple KOs), Rab11 (Rab11a/b double KOs), Rab25, and Rab35 MDCK KO cells (n=15-38 cysts per condition).

(B) PAP stages (24h) of MDCK WT and Rab3, Rab11, Rab25 and Rab35 MDCK KO cells. Cells were stained with anti-ezrin and anti-PatJ antibodies, and with phalloidin.

(C) MDCK cells were cultured in Matrigel in the presence of the indicated inhibitors/activators. Cells were fixed at 48h and stained with anti-ezrin and anti-CGN antibodies. Representative 2-cell (PAP) stages are shown.
(D) MDCK cells were cultured in Matrigel in the presence of the indicated inhibitors/activators. Cells were fixed at 48h and stained with anti-ezrin and anti-CGN antibodies. Representative 3-4-cell stages are shown.
(E) MDCK cysts were treated with CNO3 (a RhoA activator) or endosidin-2 (an exocyst inhibitor) for 72h, fixed and stained with anti-PODXL antibodies.
(F and G) Quantification of lumen circularity and lumen phenotype based on data shown in (C-E). * $p<0.05$; ** $p<0.01$; Student's t-test (n=12-21 cysts).
