## Supplementary File S2 for "Bulk delivery of a preassembled apical surface initiates epithelial lumen formation"

**Supplemental Table 1:**

| Antibodies | | |
| --- | --- | --- |
| Mouse monoclonal anti-Pals1 (G-5) | Santa Cruz | sc-365411  RRID: AB_10851475 |
| Rabbit polyclonal anti-PatJ | LSBio | LS-C410011 |
| Mouse monoclonal anti-Ezrin (3C12) | Invitrogen | MA5-13862  RRID: AB_10979020 |
| Mouse monoclonal anti-Podocalyxin | DSHB | 3F2/D8  RRID: AB_2618385 |
| Mouse monoclonal anti ZO-1 (ZO1-1A12) | Thermo Fisher Scientific | Cat# 33-9100  RRID: AB_2533147 |
| Rabbit polyclonal anti-Partitioning-defective 3 (Par3) | Merck Millipore | Cat# 07-330  RRID: AB_2101325 |
| Rabbit polyclonal anti-HOMER1 | Invitrogen | PA5-21487  RRID: AB_11155843 |
| Rabbit polyclonal anti-HOMER2 | Atlas antibodies | HPA040134  RRID: AB_10795272 |
| Mouse monoclonal anti-GAPDH | Invitrogen | Cat# sc-47724  RRID: AB_627678 |
| anti-Mouse IgG (H+L) Alexa Fluor 488 | Invitrogen | Cat# A-21202  RRID: AB_141607 |
| anti-Mouse IgG (H+L) Alexa Fluor 555 | Invitrogen | Cat# A-31570  RRID: AB_2536180 |
| anti-Mouse IgG (H+L) Alexa Fluor 633 | Invitrogen | Cat# A-21052  RRID: AB_2535719 |
| anti-Rabbit IgG (H+L) Alexa Fluor 488 | Invitrogen | Cat# A-21206  RRID: AB_2535792 |
| anti-Rabbit IgG (H+L) Alexa Fluor 555 | Invitrogen | Cat# A-31572  RRID: AB_162543 |
| anti-Rabbit IgG (H+L) Alexa Fluor 633 | Invitrogen | Cat# A-21071  RRID: AB_141419 |
| anti-rabbit IgG (H+L) HRP conjugate | Invitrogen | Cat# A16104  RRID: AB_2534776 |
| anti-mouse IgG (H+L) HRP conjugate | Invitrogen | Cat# A16072  RRID: AB_2534745 |
| DAPI | Sigma-Aldrich | Cat# D9542 |
| Phalloidin–Atto 565 | Sigma-Aldrich | Cat# 94072 |
| Rabbit polyclonal anti-L27 (PatJ) | Kind gift from Andre Le Bivic | N/A |
| Rat polyclonal anti-ZO1 | Kind gift from Sandra Citi | N/A |
| Rabbit polyclonal anti-cingulin (CGN) C532 | Kind gift from Sandra Citi. Cardellini et al., 1996 | N/A |
| FlexAble 2.0 Coralite Plus Antibody Labelling Kit | Proteintech | Cat# KFA521, KFA523 |
| Chemicals, peptides, and recombinant proteins | | |
| Paraformaldehyde | Sigma-Aldrich | Cat# 158127 |
| Biotin phenol | Iris Biotech | Cat# LS-3500 |
| Trolox | Sigma-Aldrich | Cat# 238813 |
| Glutaraldehyde (32%) | Electron Microscopy Sciences | Cat# 16220 |
| Formaldehyde (32%) EM Grade | VWR | Cat# 100504-858 |
| Diaminobenzidine (DAB) (Free-Base) | Sigma-Aldrich | Cat# D8001 |
| Durcupan ACM resin | Sigma-Aldrich | Cat# 44610 |
| Osmium Tetroxide (2%) | Electron Microscopy Sciences | Cat# 19152 |
| Potassium Ferricyanide | Sigma-Aldrich | Cat# 702587 |
| Low Molecular Weight Tannic Acid | Electron Microscopy Sciences | Cat# 21700 |
| Uranyl Acetate | Electron Microscopy Sciences | Cat# 22400 |
| Lead Citrate Trihydrate | Electron Microscopy Sciences | Cat# 17800 |
| Sequencing Grade Modified Trypsin | Promega Corporation | Cat# V5111 |
| Lysyl Endopeptidase, MS Grade | FUJIFILM Wako Pure Chemical Corporation | Cat# 125-05061 |
| cOmplete™, EDTA-free Protease Inhibitor Cocktail | Roche | Cat# 11873580001 |
| Pierce™ Protease Inhibitor Tablets, EDTA-free | Thermo Fisher Scientific | Cat# A32965 |
| Thermo Scientific™ Pierce™ Streptavidin Magnetic Beads | Thermo Fisher Scientific | Cat# 10615204 |
| EZQ™ Protein Quantitation Kit | Thermo Fisher Scientific | Cat# R33201 |
| Polyethylenimine, Linear, MW 25000, Transfection Grade` | Polysciences | Cat# 23966-2 |
| Lipofectamine 3000 Reagent | Thermo Fisher Scientific | Cat# L3000015 |
| Blebbistatin | Sigma-Aldrich | Cat# 203391 |
| Rhosin | MedChemExpress | Cat# HY-12646 |
| Y-27632 | MedChemExpress | Cat# HY-10071 |
| Endosidin-2 | MedChemExpress | Cat# HY-120821 |
| CNO3 (RhoA activator II) | Cytoskeleton | Cat# CNO3 |
| DMEM, high glucose, no glutamine, no calcium | Gibco | Cat# 21068028 |
| VECTASHIELD® Antifade Mounting Medium | Vector Laboratories | Cat# H-1000-10 |
| Bacterial and virus strains | | |
| BL21(DE3) Competent *E. coli* | NEB | Cat# C2527H |
| NEB^®^ 10-beta Competent *E. coli* (High Efficiency) | NEB | Cat# C3019H |
| Experimental models: Cell lines | | |
| MDCK-II | ATCC | CCLV Cat# CCLV-RIE 1061  RRID: CVCL_0424 |
| MDCK-II PatJ KO | This study | N/A |
| MDCK-II Pals1 KO | Groh et al., 2024 | N/A |
| MDCK-II Pals1-EGFP | Groh et al., 2024 | N/A |
| MDCK-II Pals1-APEX2-EGFP | Tan et al., 2020 | N/A |
| MDCK-II mNeonGreen-ZO1 | Pombo-Garcia et al., 2024 | N/A |
| MDCK-II Rab3a/b/c/d KO | RIKEN BRC | Cat# RCB5102 |
| MDCK-II Rab11a/b KO | RIKEN BRC | Cat# RCB5112 |
| MDCK-II Rab25 KO | RIKEN BRC | Cat# RCB5125 |
| MDCK-II Rab35 KO | RIKEN BRC | Cat# RCB5134 |
| Oligonucleotides | | |
| PatJ sgRNA1 UACACCUAACUCUAGUUCGA  PatJ sgRNA2 AAAGUUGGUCGAACAAUCUG  PatJ sgRNA3 UUAAUGGUGUCCAACUGUAU | IDT DNA | https://sg.idtdna.com/page/products/custom-dna-rna |
| Recombinant DNA | | |
| Pals1-APEX2-EGFP (Pals1-A2E) | N/A | Tan et al., 2020 |
| C1 EGFP-APEX2 | N/A | Tan et al., 2020 |
| N1 APEX2-EGFP | N/A | Tan et al., 2020 |
| LifeAct-mCherry | Addgene | #193300 |
| pEGFP C1 | Addgene | #13031 |
| pmCherry C1 | Clontech | Cat# 632524 |
| mCherry-PatJ | Pombo-Garcia et al., 2024 | N/A |
| mCherry-PatJΔPDZ6 | Pombo-Garcia et al., 2024 | N/A |
| mCherry-L27-PDZ1 | Pombo-Garcia et al., 2024 | N/A |
| mCherry-L27-PDZ1-PDZ6 | Pombo-Garcia et al., 2024 | N/A |
| mCherry-L27-PDZ1-ZO1 | Pombo-Garcia et al., 2024 | N/A |
| GFP-Crb3 | Klinkert et al., 2016 | N/A |
| Software and algorithms | | |
| ImageJ | N/A | https://Imagej.nih.gov/ij |
| MaxQuant (version 1.6.7.0) | N/A | https://www.maxquant.org/ |
| Uniprot | N/A | https://www.uniprot.org/ |
| ProTIGY | Broad Institute, Proteomics Platform | https://github.com/broadinstitute/protigy |
| Cytoscape | N/A | https://cytoscape.org |
| BioGRID | N/A | https://thebiogrid.org |
| STRING | N/A | https://string-db.org |
| Inkscape | N/A | https://inkscape.org/ |
| R (version 4.0.3) | N/A | https://www.r-project.org/ |
| Adobe InDesign | Adobe | https://www.adobe.com/sg/products/indesign.html |
| Blender v4.3.4 | Blender Development Team | https://www.blender.org |
| Phylo | Biomni Lab | https://biomni.phylo.bio |
| Other | | |
| Matrigel | Corning | Cat# 354230 |
| Transwell filter inserts (6.5 mm) | Corning | Cat# 3413 |
| Glass coverslips (12 mm, #1.5) | Electron Microscopy Sciences (EMS) | Cat# 72230 |
